# Controlling one’s world: identification of sub-regions of primate PFC underlying goal-directed behavior

**DOI:** 10.1101/2021.01.13.426375

**Authors:** Lisa Y. Duan, Nicole K. Horst, Stacey A.W. Cranmore, Naotaka Horiguchi, Rudolf N. Cardinal, Angela C. Roberts, Trevor W. Robbins

## Abstract

Impaired detection of causal relationships between actions and their outcomes can lead to maladaptive behavior. However, causal roles of specific prefrontal cortex (PFC) sub-regions and the caudate nucleus in mediating such relationships in primates are unclear. We inactivated and over-activated five PFC sub-regions, reversibly and pharmacologically: areas 24 (perigenual anterior cingulate cortex), 32 (medial PFC), 11 (anterior orbitofrontal cortex, OFC), 14 (rostral ventromedial PFC/medial OFC) and 14-25 (caudal ventromedial PFC), and the anteromedial caudate, to examine their role in expressing learned action-outcome contingencies using a contingency degradation paradigm in marmosets. Area 24 or caudate inactivation impaired the response to contingency change, while area 11 inactivation enhanced it, and inactivation of areas 14, 32 or 14-25 had no effect. Over-activation of areas 11 and 24 impaired this response. These findings demonstrate distinct roles of PFC sub-regions in goal-directed behavior and illuminate the candidate neurobehavioral substrates of psychiatric disorders including obsessive-compulsive disorder.

## Introduction

In everyday life, we continually make decisions based on our goals or go onto “autopilot” to get through the day. Normal behaviors can either be goal-directed, when performing an action to obtain a specific goal, or habitual, when a stimulus can trigger a well-learned response regardless of its consequences. The goal-directed system is needed to adapt and remain flexible to changing environments and goals, whereas the habit system reduces cognitive load. However, an excessively dominant habit system can be maladaptive in some circumstances. Problems in the co-ordination and competition between the goal-directed and habitual systems are seen both in health (Balleine and O’Doherty, 2010; de Wit and Dickinson, 2009; Dolan and Dayan, 2013; Verhoeven and de Wit, 2018) and also in neuropsychiatric disorders such as obsessive-compulsive disorder (OCD) (Gillan and Robbins, 2014; Robbins et al., 2019) and addiction (Ersche et al., 2016; Everitt and Robbins, 2005). Thus, understanding the neurobiological basis of goal-directed behaviors will provide insight into the etiology of such disorders.

To choose optimally, one needs to predict or believe that one’s action will cause the desired outcome (de Wit and Dickinson, 2009; Heyes and Dickinson, 1990). Whether or not an outcome (O) is contingent upon an action (A) depends not only on the probability (P) of the outcome occurring following the action [P(A|O)] but also on the probability of the outcome occurring in the absence of that action [P (A|¬O)]. One’s sensitivity to changes in action-outcome (A-O) contingencies can be measured using a test of contingency degradation (Balleine and Dickinson, 1998; Dickinson and Weiskrantz, 1985; Hammond, 1980; Rescorla, 1966). Persistent responding following degradation of A-O contingencies implies residual habitual control (Balleine and O’Doherty, 2010). Contingency degradation assesses beliefs about the causal nature of contingencies and complements the use of ‘outcome devaluation’ (which tests whether the desire for a goal drives actions) to measure goal-directed behavior (Heyes and Dickinson, 1990).

Evidence from human, non-human primate and rodent studies has identified candidate neural systems for goal-directed behavior in mediating A-O contingencies within sub-regions of the prefrontal cortex (PFC), including ventromedial PFC (vmPFC), medial PFC (mPFC), orbitofrontal cortex (OFC), and the anterior cingulate cortex (ACC), as well as the caudate nucleus subcortically (reviewed in Balleine and O’Doherty 2010). In human functional magnetic resonance imaging (fMRI) studies, sectors of the vmPFC and anterior caudate were more active when the subjects’ actions were highly predictive of the outcome than when their actions did not predict the outcome well [that is, relating to P(A|O)] (Liljeholm et al., 2011; Tanaka et al., 2008). Studies of humans with vmPFC damage reveal intact learning of A-O contingencies but reduced awareness of such relationships under certain conditions (O’Callaghan et al., 2019; Reber et al., 2017). The vmPFC, however, is a large, heterogeneous region (Roberts and Clarke, 2019) comprising a number of cytoarchitectonically (and likely functionally) distinct regions that cannot easily be resolved with fMRI and cannot be distinguished in human lesion studies, where lesions extend across the vmPFC and do not respect architectonic boundaries.

Localized intervention studies in non-human animals have shed light on the causal role of PFC sub-regions in goal-directed actions. In non-human primates, non-specific ablations of dorsal ACC in macaques impaired their ability to adapt their actions to changes in outcome probabilities (Chudasama et al., 2013; Kennerley et al., 2006; Rudebeck et al., 2008). More selective excitotoxic lesions of area 32 (mPFC) or area 11 (anterior OFC; antOFC) in marmosets impaired the initial acquisition of A-O contingencies, reflected in their subsequent insensitivity to contingency degradation (Jackson et al., 2016). Similar insensitivity has been described in rats with excitotoxic lesions of the prelimbic cortex (PL; variously considered mPFC/vmPFC), lateral OFC and the posterior dorsomedial striatum (Balleine and Dickinson, 1998; Corbit and Balleine, 2003; Ostlund and Balleine, 2007; Yin et al., 2005). In contrast, anterior medial OFC lesions had no effect on this task, whilst affecting outcome devaluation (Bradfield et al., 2015). Thus, while it is clear that altered activity across a number of regions within prefrontal and cingulate cortices in humans and non-human animals can affect goal-directed behavior, differences in the test paradigms used and the extent of cross-species homology in prefrontal function hamper translation of the findings (Laubach et al., 2018; Roberts, 2020). For example, recent studies of vmPFC function in marmosets have highlighted functional discrepancies between this region in rodents versus primates (Roberts and Clarke, 2019; Wallis et al., 2017). Consequently, the present study provides a comprehensive comparison of the causal contribution of the perigenual ACC, vmPFC and OFC to goal-directed behavior, as measured by the sensitivity to contingency degradation in a New World primate, the common marmoset.

The structural organization of the marmoset PFC bears a greater resemblance to that of humans than to rodents (Burman and Rosa, 2009; Roberts et al., 2007; Vogt et al., 2013) and hence provides an important bridge between rodent and human studies. Moreover, the animal studies cited above mainly used contingency degradation to assess whether initial learning was goal-directed or habit-based, but not in the subjects’ ability to respond to changes in A-O contingencies in established goal-directed behavior. We modified the classic rodent task (Balleine and Dickinson, 1998) and employed a within-subjects design to ensure that animals were already exhibiting goal-directed actions prior to repeated acute manipulations in multiple brain regions. We examined the contribution of five prefrontal and cingulate sub-regions: areas 32 (mPFC), 24 (perigenual ACC), the boundary between area 14 and 25 (area 14-25; caudal vmPFC), 11 (antOFC), and 14 (rostral vmPFC/mOFC) using temporary pharmacological inactivation via local microinfusion of a combination of a GABA_A_ agonist (muscimol) and a GABA_B_ agonist (baclofen) (“mus/bac”). As inactivation of pre-genual area 24 disrupted the sensitivity to contingency degradation and this region sends dense projections into the anteromedial sector of the caudate nucleus, we inactivated the caudate, using the glutamate receptor antagonist CNQX. Because OFC and ACC were overactive in obsessive-compulsive disorder (OCD) patients, which might underlie the deficits in goal-directed behavior seen in this disorder (Robbins et al., 2019) including impaired knowledge of A-O contingencies in contingency degradation (Vaghi et al., 2019), we also determined the effects of over-activation of prefrontal areas in marmosets. We achieved this via a glutamate reuptake inhibitor, dihydrokainic acid (DHK), which increases the extracellular levels of glutamate and enhances excitability of cortical areas (Alexander et al., 2019; Bechtholt-Gompf et al., 2010; Munoz et al., 1987). In all cases, infusions were performed during contingency degradation probe sessions as well as during baseline sessions.

## Results

### Novel procedure established marmosets’ sensitivity to contingency degradation

Prior to intra-cerebral infusions, we established that marmosets behaved in a goal-directed manner (Figure 1A). Marmosets were trained to associate each of two actions with a different outcome (juice rewards; Figure 1B). In the two consecutive probe sessions in which the action-outcome (A-O) contingencies were modified, they reduced their responding when one of the two A-O associations was degraded (i.e., when “free” and response-contingent rewards were the same juice) compared with when it was not degraded (i.e., when free and response-contingent rewards were different; Figure 1D). Importantly, marmosets did not alter their responding when the A-O associations were not degraded (alternate reward presented freely) when compared with sessions where there were no free rewards (Figure 1D). Together, this indicated that the decrease in marmosets’ responding was not simply due to the presence of free rewards, but because of the weakening of the perceived causality between a specific action and outcome.

**Figure 1.**
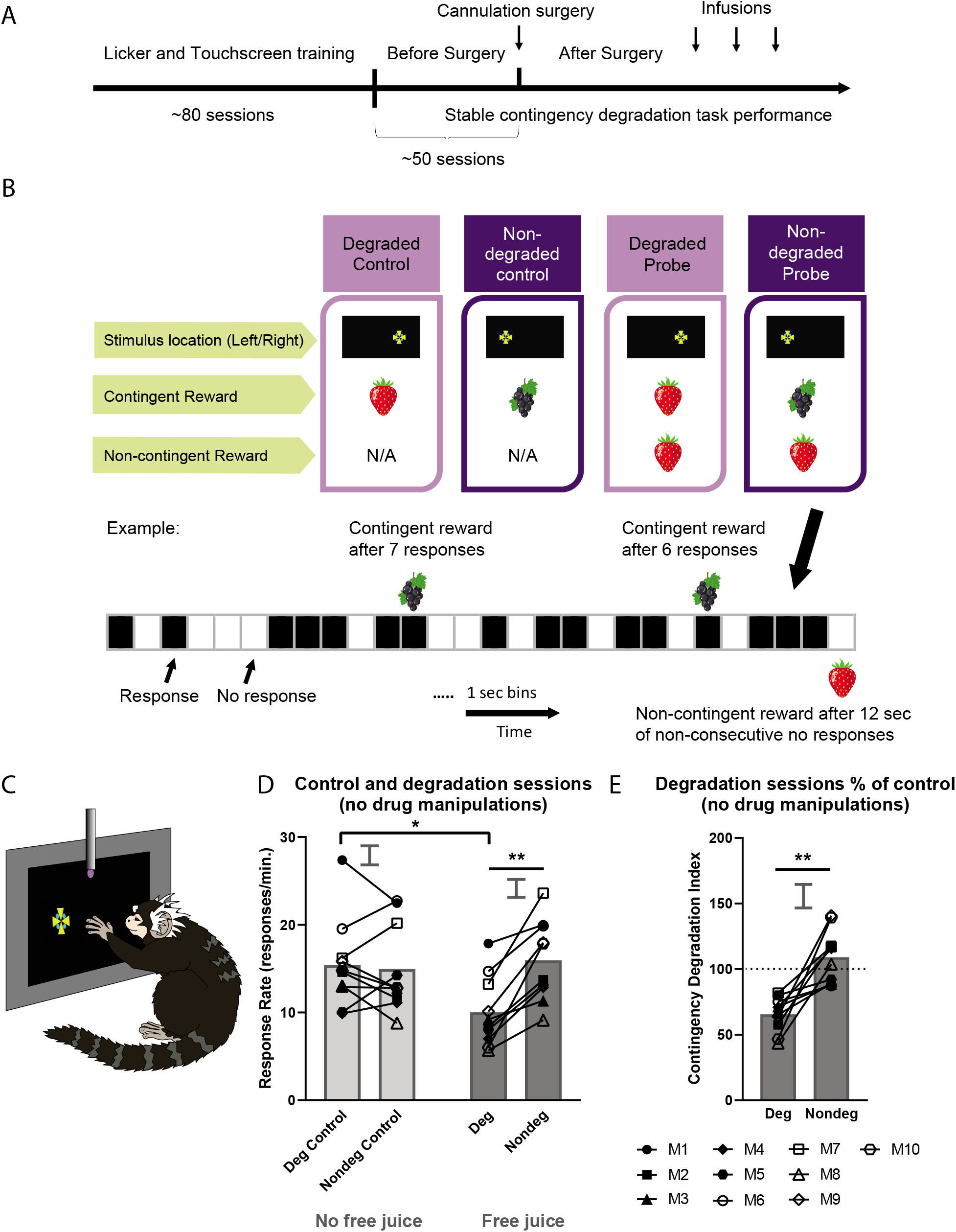
Experimental outline and a novel procedure to establish marmosets’ sensitivity to contingency degradation. **(A)** Timeline of the experiment. Marmosets were pre-trained to engage with the testing apparatus and the reward being delivered from the licking spout before being given touchscreen training (see Methods for more detail). The first drug manipulation was conducted after sensitivity to contingency degradation had been established. **(B)** The novel contingency degradation task. The first two days were control sessions in which animals responded on variable ratio schedules for each of two rewards, one of which would be degraded in the subsequent probe sessions (degraded control) and one of which would not (non-degraded control). The subsequent two days were the contingency degradation probe sessions. In this figure, the example of degraded reward was strawberry juice and the non-degraded reward was blackcurrant juice. In the degraded probe session, the response-contingent reward (strawberry juice; example reward delivery probability = 0.1) was the same as the response-non-contingent, “free” reward (strawberry juice; example probability = 0.067). In the non-degraded probe session, the response-contingent reward (blackcurrant juice; example probability = 0.1) was not the same as the response-non-contingent, free reward (strawberry juice; example probability = 0.067). Delivery probability was determined by dividing the 12-minute session into 1-second bins. Black boxes indicate a response and white boxes a non-response within that 1-second bin. See Methods for more details. **(C)** The marmoset touches the Maltese cross stimulus on the left and the associated juice reward can be retrieved from the licking spout situated in front of the touchscreen, according to a pre-programmed delivery schedule and session type. **(D)** The presence of free reward only affected marmosets’ responding when it was the same as the contingent reward, i.e. when the action-outcome (A-O) contingency was degraded (free juice presence x degradation: F1, 4 = 12.744, p = 0.0234). No difference in response rate was observed between the degraded control and non-degraded control (absence of free juice; p = 0.971). There was also a significant reduction in response rate in the degraded sessions (presence of free juice) when compared to degraded controls (absence of free juice; p < 0.0001) and non-degraded controls (absence of free juice; p = 0.001). There was no significant difference when comparing non-degraded sessions with degraded controls (p = 0.954) and non-degraded controls (p = 0.677). **(E)** Marmosets were sensitive to changes in A-O contingencies. The behavioral measure was the contingency degradation index, which is the percentage of the response rate in probe compared to control sessions (see Methods). Marmosets were goal-directed in that they showed a decrease in responding to the degraded session compared with the non-degraded session (p = 0.0009). Relevant graphs show the standard error of the differences between means (2 x SED) for “degraded v. non-degraded” comparisons, appropriate for post hoc pairwise comparisons. Deg: degraded. Nondeg: non-degraded. M: Monkey. **: p < 0.01, ***: p< 0.001.

To better assess sensitivity to contingency degradation, we developed a contingency degradation index (CDI) that considered the marked individual variability in response rates shown by marmosets e.g., see Figure 1D. The CDI was defined as the percentage of marmosets’ response rate in degradation probe sessions compared to control sessions (Figure 1E; see Methods).

Marmosets (n = 10) then received drug manipulations into their respective cannulated brain regions (Table 1; see Table S3 for drugs). The cannulae placements in the PFC sub-regions and the caudate nucleus are presented in Figure 2 (see Table S2 for cannulation coordinates).

**Table 1.** Cannulation Locations for Each Marmoset. M: Monkey. A tick indicated the brain region(s) that were targeted in that particular marmoset. Italic numbers in brackets next to ticks are the number of infusions in each brain region, used to generate the data in this paper. ^: n = 5 available, n = 4 collected data. *: Area 14-25 and 14 could be reached via extending the injectors through area 24 and area 32 vertical implants, respectively.

|  | Symbol | Area 11 | Area 24 | Area 14-25* | Area 32 | Area 14* | Caudate Nucleus |
| --- | --- | --- | --- | --- | --- | --- | --- |
| Subject |  | n = 4 | n = 4 | n = 3 | n = 5^ | n = 5^ | n = 5^ |
| M1 | ● | √ (12) |  |  |  |  |  |
| M2 | ■ | √ (12) | √ (12) | √ (12) |  |  |  |
| M3 | ▲ | √ (8) | √ (12) | √ (12) |  |  |  |
| M4 | ◆ | √ (12) | √ (12) | √ (12) |  |  |  |
| M5 | ⬢ |  | √ (12) |  |  |  |  |
| M6 | ○ |  |  |  | √ (12) | √ (12) | √ (8) |
| M7 | □ |  |  |  | √ (12) | √ (12) | √ (8) |
| M8 | △ |  |  |  | √ (12) | √ (8) | √ (0) |
| M9 | ◇ |  |  |  | √ (12) | √ (12) | √ (8) |
| M10 | ⬡ |  |  |  | √ (0) | √ (0) | √ (8) |

**Figure 2.**
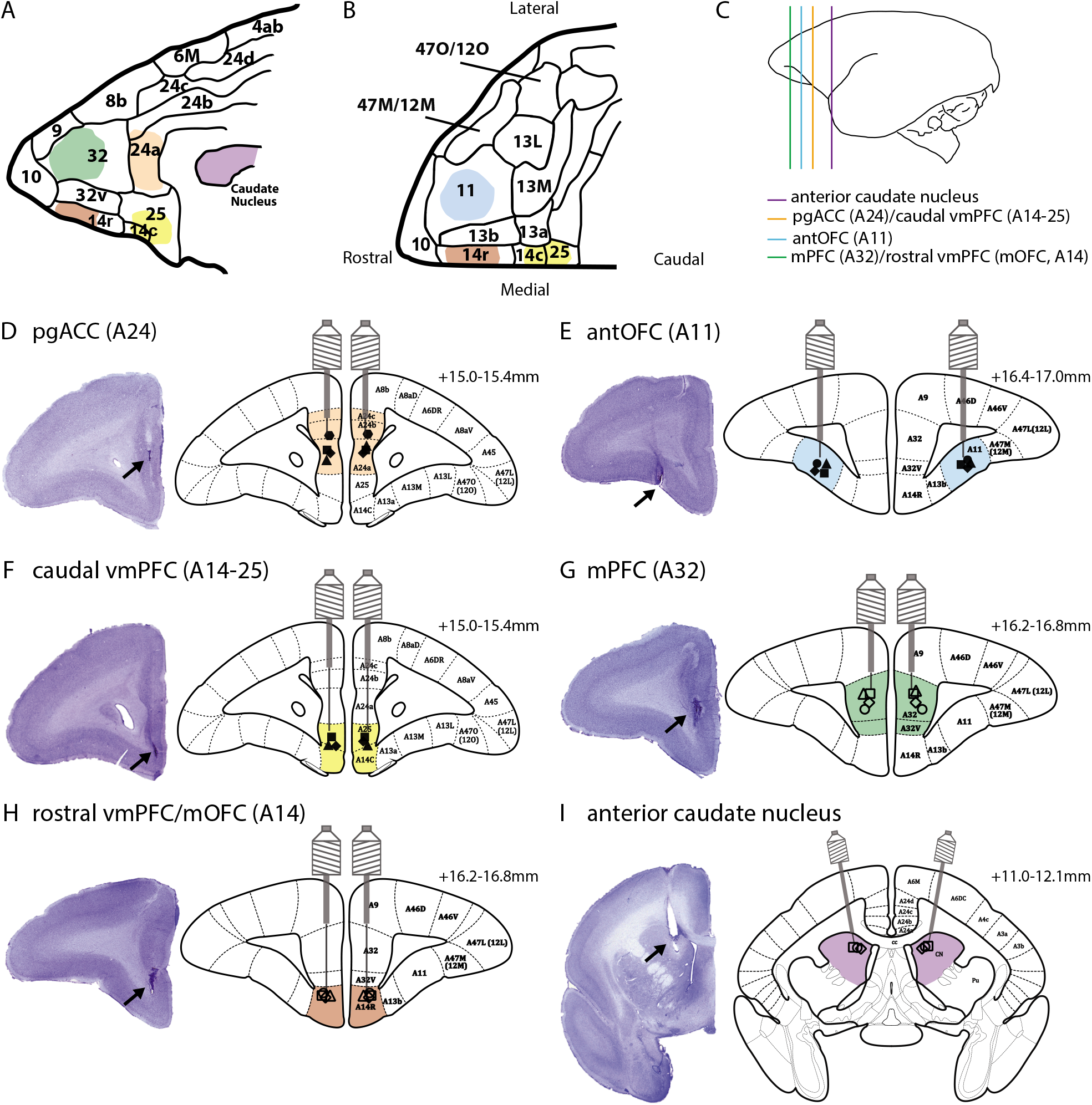
Schematic diagrams of cannulae placements in the PFC sub-regions and the caudate nucleus. **(A)** Sagittal view of the medial surface of the PFC. Each color corresponds to a targeted brain region. **(B)** Ventral view of the OFC. **(C)** Target locations of pgACC (area 24), caudal vmPFC (area 14-25), antOFC (area 11), mPFC (area 32), rostral vmPFC/mOFC (area 14) and anterior caudate in relation to the whole brain. **(D-F)** Cannulae placements in areas 11, 24 and 14-25. Area 14-25 was reached by vertically extending the area 24 injector, thus targeting both areas 24 and 14-25 via the same guide cannula. **(G-I)** Cannulae placements in areas 32, 14 and anterior caudate. Area 14 was reached by vertically extending the area 32 injector, thus targeting both areas 32 and 14 via the same cannula guide. Parcellation maps have been labeled based on Paxinos et al. (2012). See Table S2 for cannulation coordinates.

### Disrupting activity in area 24 abolished the sensitivity of actions to contingency degradation

Following either inactivation or over-activation of area 24, marmosets’ actions were insensitive to a degradation in A-O contingency (Figure 3A). Marmosets no longer distinguished between a reward that could only be obtained by performing an action (as in the non-degraded session) and one that could be obtained with or without an action (as in the degraded session). Thus, their responding was the same regardless of whether the reward could be obtained freely or not. Although both inactivation and over-activation of area 24 blunted marmosets’ sensitivity to contingency degradation, their effects on responding differed: only the effects of inactivation were specific to contingency degradation. Over-activation of area 24 made the marmosets respond less, regardless of whether the A-O contingencies were degraded; this was likely due to DHK effects that were non-specific to contingency degradation, as such effects were also seen in baseline sessions where no free reward was given (see Methods for descriptions of baseline sessions and Figure 5A).

**Figure 3.**
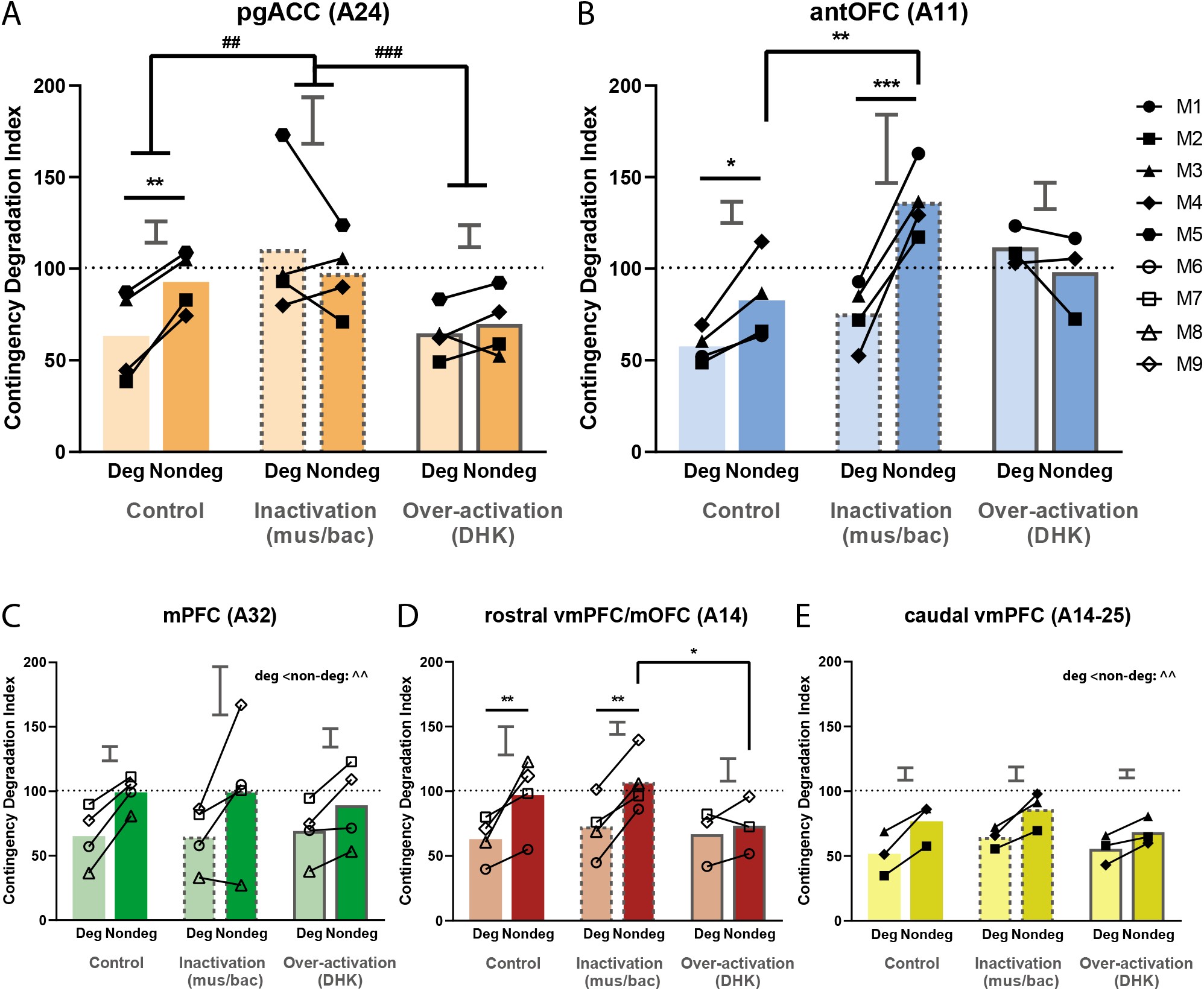
Effects of inactivation or over-activation of specific PFC sub-regions during contingency degradation. **(A)** In area 24, inactivation (via GABAA and GABAB agonism) and over-activation (via inhibition of glutamate reuptake) blunted marmosets’ sensivity to contingency degradation (treatment x degradation: F2,15 = 4.429, p = 0.0308). There was a significant difference between degraded and non-degraded sessions only following saline infusion (p = 0.0065) but not after inactivation (p = 0.331) or over-activation (p = 0.601). This lack of difference after inactivation occurred due to a selective increase in responding in degraded sessions when compared to saline (p = 0.001), whereas there was no change of responding in non-degraded sessions compared to saline (p = 0.912). Responding across degraded and non-degraded sessions following over-activation was less than that of inactivation (p = 0.0005), due to a non-specific drug effect after over-activation (see Figure 5A). **(B)** Area 11 (antOFC) inactivation apparently enhanced marmosets’ sensivity to contingency degradation, while over-activation impaired it (treatment x degradation: F2,13.287 = 7.213, p = 0.00757). Marmosets’ responding in degraded sessions was significantly reduced, compared to non-degraded sessions, under both saline (p = 0.0407) and inactivation infusions (p = 0.0004). In contrast, over-activation of area 11 abolished the degradation effect (i.e. no difference between responding in the degraded versus the non-degraded sessions; p = 0.363). Further analysis revealed a significant increase in the difference in responding between degraded and non-degraded conditions after inactivation when compared to saline infusion (p = 0.0158). This effect was driven by a significant increase in responding in the non-degraded condition after inactivation when compared to saline (p = 0.0032) but not in the degraded condition (p = 0.248). **(C)** In area 32 (mPFC), marmosets’ responding in non-degraded sessions was significantly greater than that of degraded sessions, across all treatment conditions (p = 0.0016). **(D)** A significant difference between degraded and non-degraded sessions was observed following saline infusion (p = 0.0011) and inactivation (p = 0.0012) of area 14 (rostral vmPFC/mOFC). Although no significant differences occurred between degraded and non-degraded sessions after over-activation (p = 0.445), this effect was most likely a non-specific drug effect (see Figure 5B). Responding during the non-degraded session after over-activation was significantly lower than that after inactivation (p = 0.0107) and trended lower than after saline (p = 0.0834). Conversely, the responding of marmosets during the degraded session after over-activation was not significantly different from that of inactivation (p = 0.848) or saline (p = 0.815). A similar pattern was observed in the baseline sessions, which tested the effects of drugs on marmosets’ responding without the presence of free rewards (see Figure 5B). **(E)** In area 14-25 (caudal vmPFC), marmosets’ responding in non-degraded sessions was significantly greater than that of degraded sessions across all drug conditions (p = 0.0016). Relevant graphs show 2 X SED for “degraded v. non-degraded” comparisons (area 24: n = 4; area 11: n = 4; area 32: n = 4; area 14: n = 4; area 14-25: n = 3). Deg: degraded session. Nondeg: non-degraded session. * indicates a significant effect of the degradation x treatment interaction, # indicates a significant effect between treatments, ^ indicates a significant effect between degradations. */#/^: p < 0.05, **/##/^^: p < 0.01, ***/###/^^^: p< 0.001.

**Figure 4.**
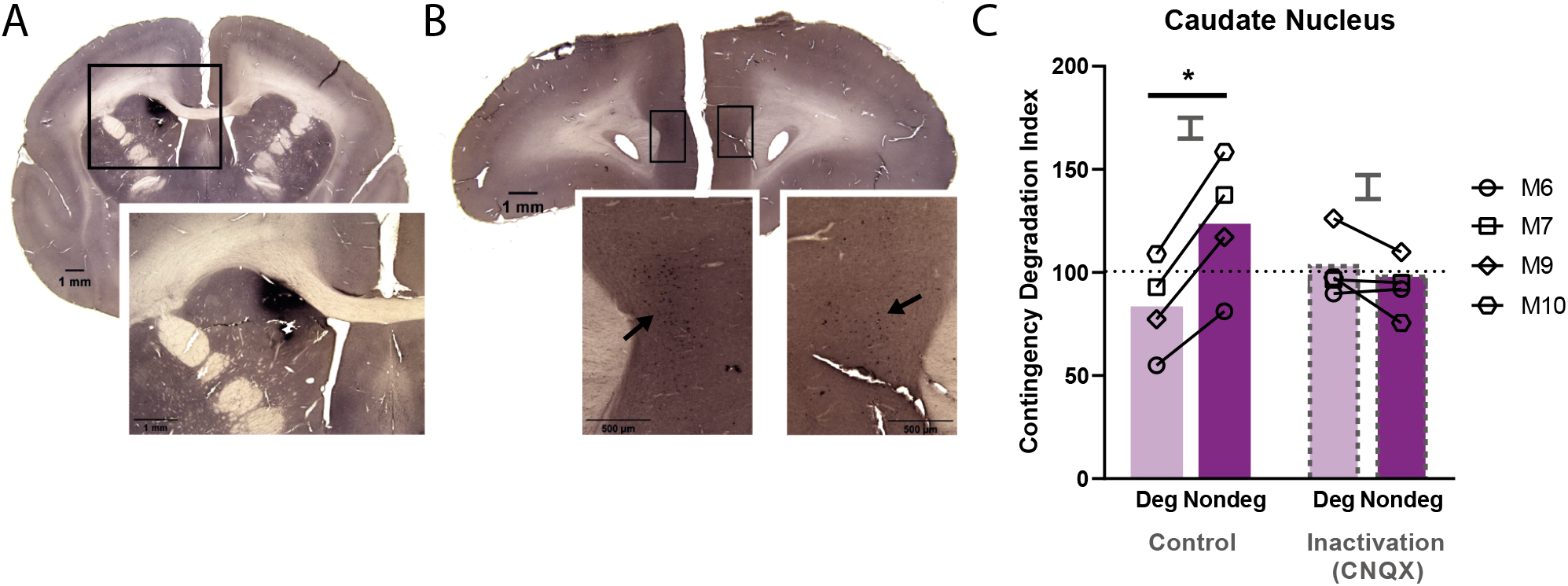
Inactivation of anterior caudate nucleus, which receives direct projection from area 24, impaired sensivity to action-outcome contingencies. **(A)** The retrograde tracer, cholera toxin B subunit, visualized in the left anterior caudate nucleus where it was injected. **(B)** Area 24, shown at the approximate placement used in this paper showing cell bodies of caudate-projecting neurons within area 24. Ipsilateral projection from area 24 to the caudate is greater than that from the contralateral projection. **(C)** Inactivation of the caudate impaired sensivity to contingency degradation. Significant treatment differences were observed on contingency degradation (treatment x degradation: F1,9 = 5.873, p = 0.0384). Inactivation (via CNQX) of the caudate nucleus that receives projection from the targeted area 24, resulted in a significant difference between degraded and non-degraded sessions following saline infusion (p = 0.0220), but not after inactivation (p = 0.523). Relevant graphs show 2 X SED for “degraded v. non-degraded” comparisons (n = 4). Deg: degraded session; Nondeg: non-degraded session. * indicates significant effect of degradation x treatment interaction. *: p < 0.05

**Figure 5.**
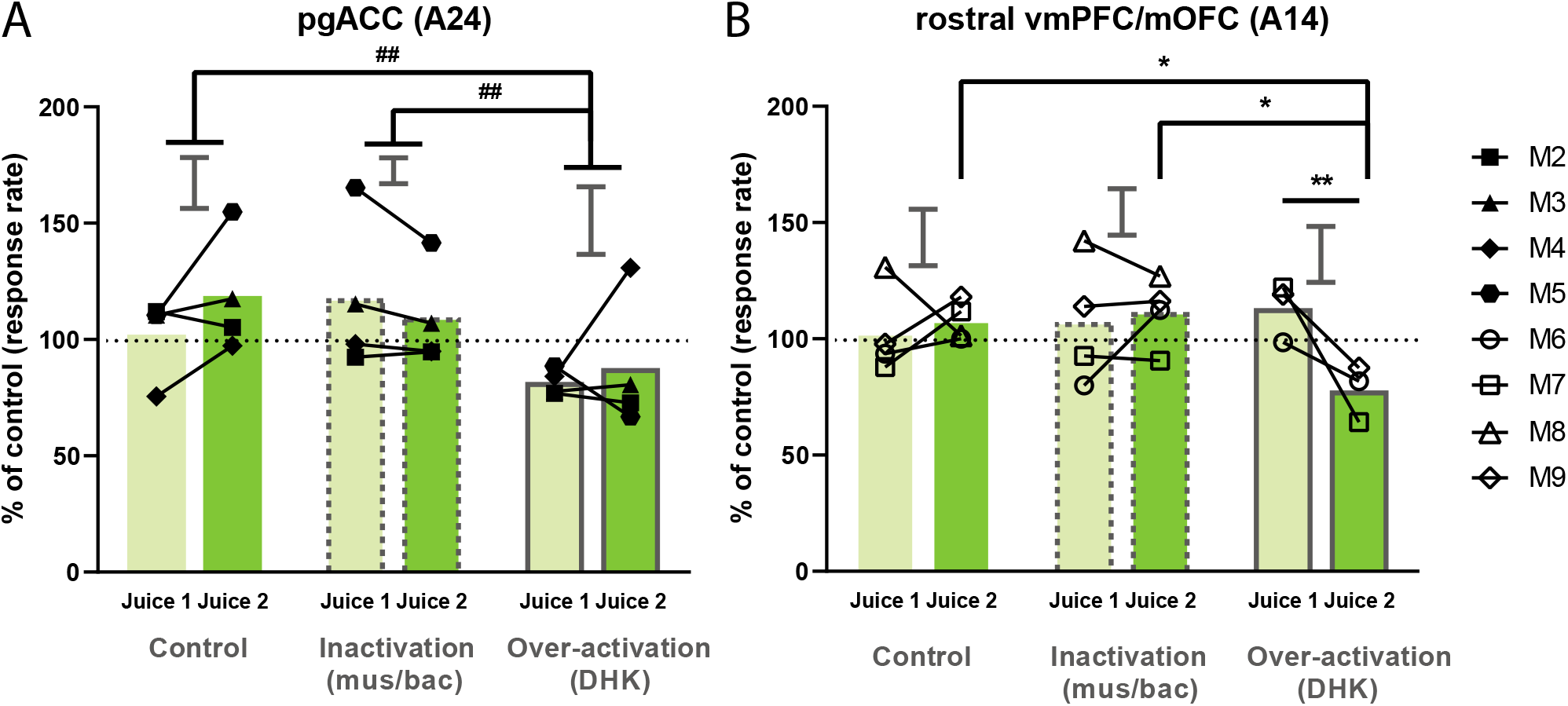
Effects of area 24 and 14 over-activation on baseline sessions. **(A)** Over-activation of area 24 significantly affected responding compared to other manipulations (treatment: F2,10 = 14.846, p = 0.00102), where it significantly decreased responding across juice 1 and 2 when compared to inactivation (p = 0.00210) or saline (p = 0.00220). **(B)** Area 14 over-activation significantly affected responding in different juice conditions (juice condition x treatment: F2,12.812 = 6.358, p = 0.0121); over-activation specifically decreased responding to juice 2, which is the contingent reward in the non-degraded session in the contingency degradation task, compared to juice 1, which is the contingent reward in the degraded session in the contingency degradation task (p = 0.0038). Responding to juice 2 after over-activation was significantly lower than that following saline (p = 0.0202) or inactivation (p = 0.0232). Conversely, responding to juice 1 after over-activation was not significantly lower than that of saline (p = 0.330) or inactivation (p = 0.556). There was no significant difference in responding after over-activation between juice 2 in baseline sessions, and the non-degraded session in the contingency degradation task (p = 0.651), whereas there was a significant difference between juice 1 and the degraded session in the contingency degradation task (p = 0.001). Relevant graphs show 2 X SED for “ Jucie 1 v. Juice 2” comparisons (area 24: n = 4; area 14: n = 4). For baseline sessions of other brain regions see Figure S2. * indicates significant effect of juice condition x treatment interaction, # indicates significant effect between treatments. */#: p < 0.05, **/##: p < 0.01

### Inactivation of area 11 enhanced, but over-activation blunted, sensitivity to contingency degradation

Inactivation of area 11 (antOFC) accentuated the difference in responding for a reward that was solely dependent on its availability through an action and one that was not (Figure 3B). This effect was driven by an increase in responding when the causal association between action and outcome was intact (non-degraded session), and not by a decrease in responding when the A-O association was weakened (degraded session). Correspondingly, the opposite effect was seen after over-activation of area 11, which blunted animals’ sensitivity towards the degradation in A-O relationships (Figure 3B). This over-activation effect was unlike that observed for area 24 because it was not accompanied by an overall decline in responding (either in the degradation test or at baseline).

### Manipulations of areas 32, 14-25, or 14 had no specific effects on the sensitivity of actions to contingency degradation

Inactivation or over-activation of either area 32 (Figure 3C) or 14-25 (Figure 3E) did not affect responsivity to contingency degradation. Similarly, inactivation of area 14 did not impair sensitivity to contingency degradation (Figure 3D). Although over-activation of area 14 did blunt the contingency degradation effect (Figure 3D), the finding that it also reduced baseline responding suggests it was not specific to contingency degradation per se (see Figure 5B). Therefore, manipulations of areas 32, 14-25 and 14 appeared not to impact, specifically, the use of previously acquired response-outcome contingencies to guide responding following contingency degradation.

### Inactivation of the anterior caudate nucleus, which receives projections from area 24, impaired sensitivity of actions to contingency degradation

After identifying area 24 as the main PFC sub-region necessary for detecting and acting upon changes in instrumental A-O contingencies (Figure 3A), we determined its target region within the caudate nucleus, as fronto-striatal pathways have been implicated in mediating goal-directed behavior (Balleine and O’Doherty 2010). Rodent and macaque tracing studies had indicated that medial PFC and dorsal ACC project to the anterior dorsal striatum (Haber et al., 1995; Ferry et al., 2000; Heilbronner et al., 2016). However, tracing studies and the Marmoset Brain Connectivity Atlas have not previously investigated the connectivity of area 24a, i.e. the pgACC targeted in the present study, with other brain regions (Majka et al., 2020; Roberts et al., 2007; also see http://www.marmosetbrain.org/). Therefore, we infused a retrograde tracer, cholera toxin B subunit, into the anterior caudate via guide cannula in a marmoset not included in the behavioral data collection (Figure 4A). Cell bodies identified in area 24 confirmed the existence of an area 24-caudate pathway (Figure 4B). Bilateral projections from area 24 were observed, with greater ipsilateral projections (Figure 4B). Besides area 24, the anterior caudate also received PFC projections from areas 8 and 32 (area 32 see Figure S3).

This region of the anterior caudate was then inactivated using CNQX (an AMPA glutamate receptor antagonist), to block excitatory glutamatergic input, including from the PFC, into this region (Galinanes et al., 2011; Darbin and Wichmann, 2008). Anterior caudate inactivation blocked the responsivity to contingency degradation (Fig. 4C).

### Regional inactivations and over-activations in baseline sessions (without degradation) had differential effects on instrumental responding

Baseline sessions were conducted separately in close temporal proximity to the contingency degradation sessions, to examine the effect of pharmacological manipulations on baseline instrumental responding. Over-activation of area 24 uniformly depressed responding when compared to saline or inactivation (Figure 5A), which mirrored the effects in the contingency degradation task (Figure 3A). Saline infusions and inactivation of area 14 did not affect responding; however, over-activation of area 14 specifically depressed responding to the response-contingent reward (Juice 2) in the non-degraded sessions of the contingency degradation task but not for the response-contingent reward (Juice 1) in the degraded sessions (Figure 5B). This specific decrease in responding may explain the decline in responding after area 14 over-activation in the non-degraded session of the main contingency degradation task (Figure 3D).

Saline, inactivation and over-activation of areas 11 or 14-25, or caudate nucleus did not alter responding during baseline sessions (Figure S2A, B, D). Across all drug manipulations in area 32 (Figure S2C), marmosets increased responding to Juice 2 compared to Juice 1; this significant effect is likely non-specific arousal associated with infusions and handling since it occurred after all infusions, including saline. Why it should be selective to Juice 2 may be related to the fact that Juice 2 was the designated preferred juice.

## Discussion

This study provides the first causal evidence in primates that area 24 (perigenual ACC) is necessary for detecting and acting on changes in instrumental action-outcome (A-O) contingencies and hence the capacity for understanding whether one’s behavior exerts control over the environment. Both inactivation and over-activation of area 24 impaired the response to contingency degradation, indicating that an optimal level or pattern of area 24 activity was required. Similar impairments were seen after reducing the excitatory input into the area of the caudate nucleus projected to by area 24, indicating the potential involvement of a fronto-striatal circuit in exerting cognitive control over voluntary goal-directed behavior. In contrast, area 11 (antOFC) appeared to have an opposing influence, with inactivation enhancing, and over-activation impairing, contingency degradation. There were no effects of inactivation or over-activation of areas 32 (mPFC) or 14-25 (caudal vmPFC) on behavior following degradation. Inactivation of 14 (rostral vmPFC/mOFC) did not affect the response to contingency degradation either, while the blunting effect seen after over-activation was likely due to a non-specific drug effect rather than insensitivity to contingency degradation. The overall findings are summarized in Table 2.

**Table 2.**
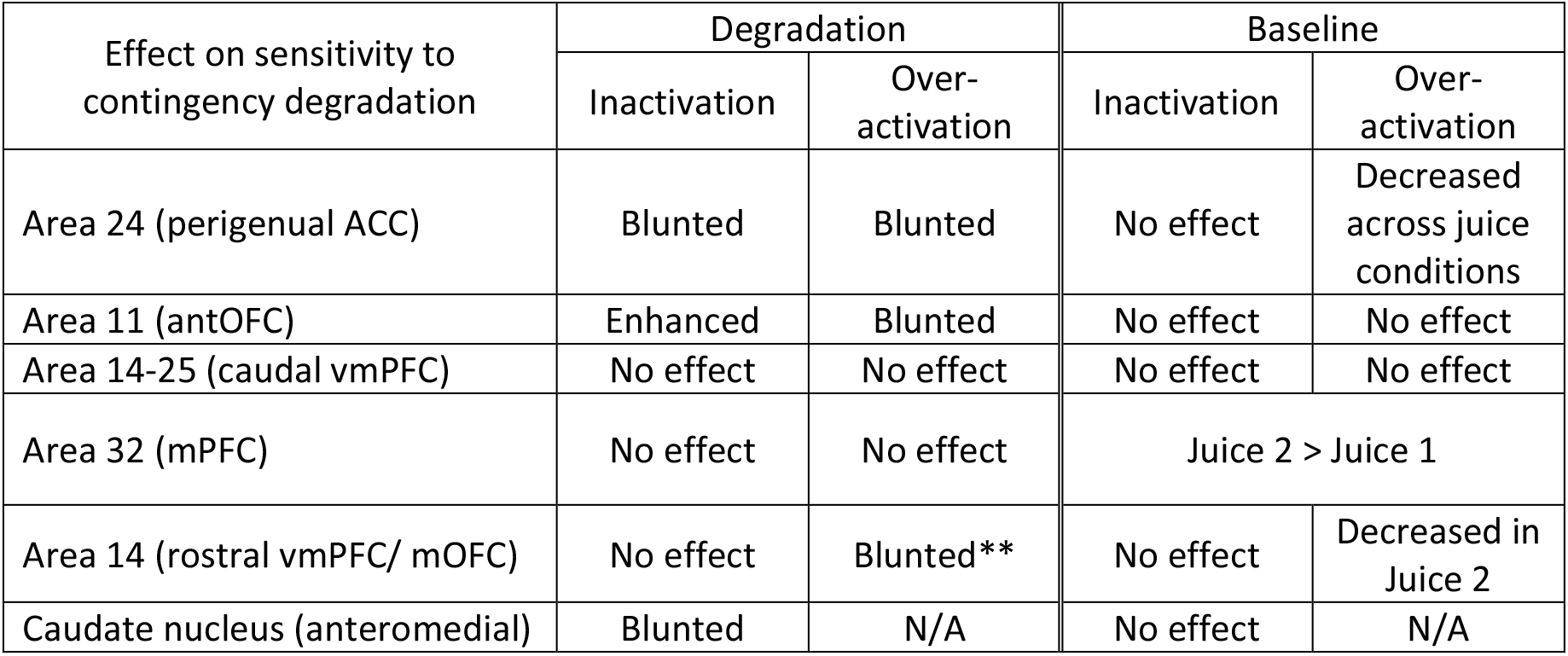
Results Summary. ** This blunting of the sensitivity to contingency degradation might be due to a non-specific drug effect observed in the baseline sessions (see Results).

The vmPFC, which has been implicated in A-O contingency learning in human neuroimaging studies, subsumes a large heterogeneous region including areas 10, 14, ventral ACC regions 25, 32, 24a, and very often in human lesion studies, orbitofrontal areas 11 and 13 (Roberts and Clarke, 2019; Schneider and Koenigs, 2017). Given this broad definition of vmPFC, it is not surprising that it has been implicated in a wide range of functions beyond A-O contingencies, including value comparison, reward processing, decision-making, threat extinction, and social cognition (Hiser and Koenigs, 2018). Our results thus define, causally, specific regions of the primate vmPFC and mPFC that mediate the detection of changed A-O contingencies, resulting in altered expression of goal-directed behavior.

### Goal-directed control over responding requires optimal levels of activity in area 24, but not area 32

Perigenual ACC (area 24) is the only PFC sub-region in this study that disrupted animals’ sensitivity to the current A-O contingencies when either inactivated or over-activated. The impaired sensitivity after area 24 over-activation was nevertheless qualitatively different from that of inactivation. After area 24 inactivation, marmosets maintained their responding in the non-degraded session but did not reduce responding in the degraded session. However, after area 24 over-activation, while animals were impaired in differentiating their responses to degraded versus non-degraded sessions, they also decreased their responding uniformly across both sessions, potentially from a generalized drug action observed on baseline sessions, in which drug infusions were made without free rewards. Nonetheless, our result is consistent with previous literature that ACC mediates reward-guided decision making (Chudasama et al., 2013; Hayden et al., 2009; Hayden and Platt, 2010; Holroyd and Coles, 2002; Holroyd and Yeung, 2012; Rushworth and Behrens, 2008; Rushworth et al., 2007; Rushworth et al., 2004; Wallis and Kennerley, 2011). Specifically, the firing rate of rostral ACC neurons tracks positive prediction error, unexpected reward delivery by the subject (Matsumoto et al., 2007), outcome surprisingness (unassigned prediction error), and likelihood of adjusting behavior (Hayden et al., 2011). The rostral ACC appears also to be important for selecting and maintaining learned task information across time and sessions (Amiez et al., 2006; Kennerley et al., 2006; Seo and Lee, 2007). Therefore, one possible explanation for the marmosets’ impairment in the contingency degradation task could be an inability to reduce their responding in the degraded condition because of a failure to track the changes in the consequences of their actions across the session.

Inactivation or over-activation of area 32 did not affect performance after contingency degradation; these findings are significant given previous rodent evidence that contingency degradation is impaired by PL lesions (Balleine and Dickinson, 1998; Corbit and Balleine, 2003). However, it is unclear how the rat PL relates to primate areas 24 and 32 (Heilbronner et al., 2016; Vogt et al., 2013), with the definition varying depending on criteria based on cytoarchitecture (Vogt et al., 2013), connectivity (Heilbronner et al., 2016) or function (Milad et al., 2007). The lack of involvement of area 32 in the current contingency degradation task stands in contrast to its role in the initial learning of A-O associations, shown via excitotoxic lesions of area 32 in the marmoset (Jackson et al., 2016). This is also consistent with evidence that excitotoxic PL lesions in rats impaired the acquisition of contingency learning (Balleine and Dickson, 1998; Corbit and Balleine, 2003), if PL is homologous or analogous to area 32. Thus, area 32 may be necessary only for acquiring goal-directed contingencies and not for their expression, a dissociation yet to be investigated in rodent PL. Nevertheless, such a dissociation is seen with respect to the sensitivity to outcome devaluation following PL lesions (Ostlund and Balleine, 2005; Tran-Tu-Yen et al., 2009). Therefore, area 24 (and not area 32) may be needed instead to express the effects of contingency degradation knowledge, highlighting a possible anterior to posterior transfer of information in the medial PFC. Indeed, Tang et al. (2019) have suggested the boundary area of pregenual area 24 and area 32 to act as a central hub for integrating information from sensory, motoric, limbic and executive decision-making regions, based on anatomical connectivity patterns in macaques and humans.

### Excitatory projections from area 24 to the caudate nucleus may affect behavioral expression of action-outcome contingencies

As predicted, inactivation of the anterior caudate blunted animals’ sensitivity to contingency degradation. This is consistent with studies showing that anterior caudate activity is involved in mediating A-O contingencies in humans (Liljeholm et al., 2011; Tanaka et al., 2008; Tricomi et al., 2004). In rats, the putative homolog of the caudate, the dorsomedial striatum (DMS), is also implicated in goal-directed behavior; in particular findings have highlighted the posterior DMS (Hart et al., 2018a; Hart et al., 2018b; Yin et al., 2005), although there is increasing evidence for the involvement of anterior DMS as well (Corbit and Janak, 2010; Shipman et al., 2019). We verified that the targeted anterior caudate in the marmoset receives projections from area 24 employing retrograde tract-tracing. As these projections are known to be glutamatergic, we used CNQX, an AMPA-receptor antagonist, to block input into the caudate, including that from area 24 as well as from other areas of the cortex and subcortical regions (Haber, 2003; Russchen et al., 1985). Thus, it can be hypothesized that a major output from area 24 for the expression of contingency degradation is to the anterior caudate. Of note, just as area 24 has been proposed as a connectivity hub (Tang et al., 2019), so too its striatal projection area, the rostral dorsal caudate, may serve to integrate inputs from other critical areas implicated in contingency knowledge (Liljeholm et al., 2011), including OFC, ACC and inferior parietal lobule (Choi et al., 2017). Given the mutual connections between areas 32 and 24, and their overlapping striatal projections to the anterior caudate (Averbeck et al., 2014; Draganski et al., 2008; Mailly et al., 2013; Roberts et al., 2007; Figure S3), future studies should investigate the functions of this putative network in controlling the acquisition and expression of instrumental A-O contingencies.

### Areas 14/14-25 are not specifically implicated in the response to contingency degradation

Areas 14 (rostral vmPFC/mOFC) and 14-25 (caudal vmPFC) were not specifically involved in response to changes in A-O contingencies. The complete lack of effects of inactivation is consistent with rodent studies, in which lesions of a putative homolog of these regions, the anterior mOFC, impaired the effects of outcome devaluation but not contingency degradation (Bradfield et al., 2015; Bradfield et al., 2018). Although marmosets receiving over-activation of area 14 did not differentiate between degraded and non-degraded sessions, the finding that baseline responding was also affected prevents any firm conclusions concerning contingency degradation. Consistent with a lack of involvement of area 14 in contingency degradation is the hypothesis that this region tracks and contrasts the intrinsic representations of action-associated outcome values during alternative choice situations (Noonan et al., 2010; Padoa-Schioppa and Assad, 2006; Rudebeck and Murray, 2011; Stalnaker et al., 2015; Valentin et al., 2007; Wallis and Kennerley, 2011). Indeed, the decline in responding after area 14 over-activation on baseline sessions is consistent with reported blunting of anticipatory arousal to high-value food reward in marmosets (Stawicka et al., 2020). Therefore, while area 24 could be important for mediating ‘causal beliefs’ about behavior, area 14 may be more critical for comparative valuations in choice. Although imaging studies (Liljeholm et al., 2011; Tanaka et al., 2008) have shown a positive correlation between objective measures of causality and blood-oxygen-level-dependent (BOLD) activity within vmPFC, it is unclear whether this region is area 10 (Price, 2007) or 14 (Mackey and Petrides, 2010).

### Inactivation of area 11 may enhance, and over-activation impair, expression of action-outcome associations, putatively due to competition between Pavlovian and instrumental systems

Much evidence supports a role for OFC in acquiring and updating new information when tasks have strong Pavlovian components in both monkeys (Murray et al., 2007; Noonan et al., 2010; Rudebeck et al., 2008; Rushworth et al., 2007; Walton et al., 2010) and rats (Balleine et al., 2011; Ostlund and Balleine, 2007; Panayi and Killcross, 2018; Parkes et al., 2017). Although OFC impairments have been observed using stimulus-reinforcement learning tasks (Murray et al., 2007; Rolls, 2004), the OFC does not appear essential for the instrumental control of behavior (Ostlund and Balleine, 2007; Rudebeck et al., 2008), though see (Gremel and Costa, 2013; Zimmermann et al., 2017). However, it has been implicated in mediating Pavlovian-to-instrumental transfer effects (Cartoni et al., 2016; Holmes et al., 2010). In the current study, inactivation of area 11 enhanced the effect of contingency degradation whereas over-activation impaired it, suggesting that this region most likely exerts interfering Pavlovian control over instrumental responding. Specifically, in the current paradigm, instrumental responding to either the left or the right side of the touchscreen according to the specific A-O association (e.g. left-blackcurrant juice; right-strawberry juice) may be subject to interference by parallel Pavlovian approach responses, since the visual stimuli associated with each reward were identical (Figure 2B, C). Thus, inactivation of area 11 may have reduced Pavlovian interference and hence enhanced instrumental performance, while over-activation produced the opposite effect (i.e. increased Pavlovian interference). In contrast, in our previous study of contingency degradation (Jackson et al., 2016), the distinct visual properties of the stimuli presented on the left or right differentially predicted the outcome and may thus have formed Pavlovian stimulus-outcome associations that facilitated performance. This may explain why OFC (area 11/13) lesions impaired contingency learning in that study. The present findings of contrasting, potentially conflicting, interactions by different PFC sub-regions mediating instrumental goal-directed behavior agrees with other recent formulations (Balleine, 2019; O’Doherty et al., 2017; Rushworth et al., 2011).

### Methodological considerations, controls and limitations

This study employed an established method for inactivating cortical areas, using intracerebral infusions of a mixture of GABA_A_ and GABA_B_ receptor agonists. The possibility of diffusion from the site of infusion is relatively slight in relation to the volume of the different PFC regions but in any case, the dissociable and selective nature of effects obtained suggests that such diffusion did not occur to any major extent. These regions were also over-activated by infusions of the astrocytic excitatory amino acid transporter 2 (EAA2/GLT-1) inhibitor DHK (Anderson and Swanson, 2000; Arriza et al., 1994), which previously has been shown to increase local concentrations of extracellular glutamate and to increase the excitability of the neuronal population and post-synaptic action as shown by microdialysis (Fallgren and Paulsen, 1996; Munoz et al., 1987), electrophysiology (Munoz et al., 1987), FDG-PET (Alexander et al., 2019) and immediate early gene *c-fos* expression (Alexander et al., 2019; Bechtholt-Gompf et al., 2010).

Baseline sessions controlled for effects of DHK on instrumental responding independent of responsivity to contingency degradation per se. We observed baseline response rate decreases for areas 24 and 14, the latter in one condition only.

### Implications

We show specific causal contributions of area 24 of the primate prefrontal cortex to the detection and expression of A-O contingency changes as part of the control of goal-directed behavior. Persistence of responding during contingency degradation has been interpreted as an expression of habitual control (Balleine and Dickinson, 1998) although this is not necessarily the case (de Wit et al., 2018; Robbins and Costa, 2017), so further studies are required to establish whether area 24 exerts control over habits, in addition to goal-directed behavior. Contingency management can also be impaired following other PFC manipulations, such as over-activation of the anterior OFC (area 11) or area 24. These findings have implications for human psychiatric disorders such as OCD and schizophrenia, both of which involve impairments in goal-directed behavior (Barch and Dowd, 2010; Gillan et al., 2014; Morris et al., 2015). A recent study (Vaghi et al., 2019) found that OCD patients over-responded when response contingencies were manipulated to degrade the A-O contingency by providing ‘free’ reinforcement as in the present study. OCD patients are known to have over-active regions of the PFC, notably the ACC and OFC (Baxter et al., 1988; Fitzgerald et al., 2011; Gillan and Robbins, 2014; Maia et al., 2008; Menzies et al., 2008; Pauls et al., 2014; Robbins et al., 2019; Whiteside et al., 2004), especially following symptom provocation (Nakao et al., 2005; Rauch et al., 1994). Our findings concerning the over-activation of both areas 24 and 11 are consistent with the pathophysiology of OCD and may indicate a possible role for maladaptive Pavlovian-to-instrumental transfer effects (Bradfield et al., 2017). Moreover, schizophrenia has been associated with a loss of GABA-ergic neurons in the anterior cingulate cortex (de Jonge et al., 2017); this might be associated with the impairments in goal-directed behavior seen in people with schizophrenia, which may underlie the ‘negative’ symptoms of schizophrenia (Morris et al., 2018).

## Conclusion

The perigenual cingulate cortex (area 24) in the marmoset monkey is identified as a key cortical region in the detection and/or expression of changes in action-outcome contingencies. Other PFC regions, including anterior OFC (area 11), rostral (area 14) and caudal vmPFC (area 14-25) and area 32 in the mPFC, appear less involved, with inactivation of area 11 actually enhancing (and over-activation impairing) sensitivity to action-outcome (A-O) contingencies. Our findings have implications for understanding the neural control of goal-directed behavior and for certain psychiatric disorders, including OCD and schizophrenia.

## Acknowledgments

The work was supported by a Senior Investigator Award from the Wellcome Trust to T.W.R. (104631/Z/14/z) and conducted in the Behavioural and Clinical Neuroscience Institute. L.Y.D. is in receipt of the Angarrad Dodds John Bursary in Mental Health and Neuropsychiatry. We thank Gemma Cockcroft and Lauren McIver for histology, and Hannah F. Clarke and Christian M. Wood for surgery.

## Author Contributions

Conceptualization, L.Y.D., N.K.H., S.A.W.C., R.N.C., A.C.R., T.W.R.; Methodology, L.Y.D., N.K.H., S. A.W.C., N.H., R.N.C., A.C.R., T.W.R.; Validation, L.Y.D., N.K.H., S. A.W.C., N.H.; Formal Analysis, L.Y.D., N.K.H., S. A.W.C., R.N.C.; Investigation, L.Y.D., N.K.H., S. A.W.C., N.H.; Resources, T.W.R., A.C.R.; Writing – Original Draft, L.Y.D., A.C.R., T.W.R.; Writing –Review & Editing, all authors; Visualization, L.Y.D. N.K.H.; Funding Acquisition, T.W.R.; Supervision, A.C.R., T.W.R., N.K.H.

## Methods

### Experimental Model and Subject Details

#### Common marmoset (Callithrix jacchus)

Ten common marmosets (*Callithrix jacchus*; four males and six females) were used for data collection for the contingency degradation task, while one marmoset (female) was used for tract-tracing. All were experimentally naïve at the start of the study. They were housed and bred on-site in a conventional barrier facility in the University of Cambridge Marmoset Breeding Colony. Experimental animals were housed in male-female pairs in custom-made housing (Tecniplast UK Ltd., Kettering, UK). The rooms were kept at a constant temperature of 24° C and relative humidity of 55%. The rooms were illuminated gradually from 7:00 am to 7:30 am and dimmed from 7:00 pm to 7:30 pm to simulate the day/night cycle. The marmosets were tested 4-5 days per week and not at the weekends. All monkeys were fed 20 g of MP. E1 primate diet (Special Diet Services) and sliced carrots on five days a week after the daily behavioral testing session, with simultaneous free access to water for two hours. On weekends, their diet was supplemented with fruit, rusk, malt loaf, eggs, bread, and treats, and they had free access to water. The male marmosets were vasectomized to prevent pregnancy of their female partners. Their home cages were filled with environmental enrichment such as ropes and ladders. All animals were carefully monitored by the unit Named Animal Care and Welfare Officer (NACWO), researchers, the Named Veterinary Surgeon (NVS), and animal technicians. The projects were conducted under Home Office Project Licenses 70/7618 and P09631465, and all studies were verified and authorized by the unit NACWO. The projects were regulated under the Animals (Scientific Procedures) Act 1986 Amendment Regulations 2012 following ethical review by the University of Cambridge Animal Welfare and Ethical Review Body (AWERB).

### Method Details

#### Behavioral testing apparatus and paradigm

##### Testing Apparatus

Testing took place using an automated touch-screen apparatus (Biotronix, Cambridge, UK). Marmosets were transferred from their home cages to the testing apparatus via a transparent Perspex box, which is designed to be inserted directly into the testing apparatus for the duration of testing. The marmoset could move freely within the box and was not otherwise restrained. One side of the box was opened to enable the marmosets to interact with computer-controlled stimuli presented on a touchscreen (Campden Instruments, Loughborough, UK). They received liquid reinforcements from a spout/licker that was suspended centrally in front of the touchscreen (Figure 1C), which could deliver up to four different liquid rewards. The experiments were monitored and could be recorded by mounted cameras in the testing chamber. The MonkeyCantab program (R.N. Cardinal) controlled the touchscreen, pumps, spout and speakers via the Whisker control system (Cardinal and Aitken, 2010) (Cardinal and Aitken 2010).

##### Licker and touchscreen training

The animals went through licker and touchscreen training before progressing to the contingency degradation task (Figure 1A). The main food reinforcer (banana milkshake, Nesquik) was initially introduced into the marmosets’ home cages and they were transferred to the testing apparatus for familiarization. They were shaped to approach the licking spout without experimenter guidance. The reward was delivered freely through the licking spout in the testing apparatus according to a fixed schedule: 8-s reward with 8-s inter-trial intervals (ITIs). During all reward delivery, an auditory cue (‘birdsong’) was also played. There were three phases of touchscreen training and each phase was completed in separate training sessions (Table S1). In the initial phase of touchscreen training, animals responded to a horizontal green bar that spanned the width of the touchscreen. Banana milkshake was delivered as a reward for 8 seconds. In the second phase, animals responded to a small green square in the center of the touchscreen. In the final training phase, the same green square was randomly presented to the left or right of the center of the touchscreen. After training on a fixed ratio 1 schedule, in which each response was reinforced, animals were switched to a variable ratio (VR) 3 schedule, in which they received a reward after every 2-4 responses. They then moved to a VR 6 schedule, and eventually a VR 10 (range 5-15 responses per reward) schedule. Following stable performance (3 consecutive sessions of consistent responding), the banana milkshake was replaced with blackcurrant, strawberry, summerfruit, or apple and mango juice. Each animal was assigned a pair of juices, with one juice always associated with the left stimulus, and the other the right stimulus. After another 3 stable sessions of performance, animals were transferred to the final contingency schedule (described in detail below) in which the green square was replaced with a compound, multi-colored stimulus (Maltese cross; Figure 1C). The sequence of touchscreen training is summarized in Table S1.

##### Contingency degradation task

The contingency degradation task measures goal-directed behavior (action-outcome associations). It used a four-day block design consisting of two control sessions followed by two contingency degradation probe sessions (Figure 1B). In the first two (control) sessions, animals responded to one of the stimuli (left or right location) for response-contingent reward in the first session, and the other stimulus on the opposite location for a different contingent reward in the second session. The two stimuli were identical and only differed in their display location (i.e. either on the left or the right of the center of the touchscreen, never displayed concurrently). Performance across the first two control sessions provided control for comparison against the subsequent two additional degradation probe sessions. In the degradation probe session, the non-contingent, ‘free’ reward was introduced. In one session, the non-contingent reward was the same as the contingent reward, resulting in contingency degradation (degraded, action-outcome association weakened). In the second session, the non-contingent reward was the alternative reward not contingently available in that session, thus maintaining action-outcome associations for the contingent reward (non-degraded).

To implement these degradation schedules, each 12-minute session was divided into 1-s bins. The mean probability of receiving the contingent reward was p = 0.1, i.e. an average of 10 responses would yield a reward (VR 10, range 5-15). Because of the large individual variance in response rate between marmosets, the probability of receiving the non-contingent reward was customized for each animal, and determined to ensure that they would detect the free rewards but not so many free rewards as to produce satiety and lead to the cessation of responding. For example, if p = 0.067, for every 1-sec bin when the animals did not respond, the mean probability of non-contingent reward delivery was 0.067 (1 reward delivered on average every 15 sec of non-responding, range 10-20 sec).

Pharmacological manipulations of the brain occurred within subjects. They received infusions on the final two sessions of the contingency degradation probe sessions (degraded and non-degraded).

Animals that are sensitive to the action-outcome contingencies will show a much greater reduction in responding when the non-contingent juice is the same as the contingent juice (degraded condition) but not when it is different (non-degraded condition). However, this differential effect is partly dependent on animals preferring to get access to a variety of rewards rather than one reward and/or preferring a total quantity of juice greater than that only provided for free. When assigning juices to marmosets we tried to use a pair of juices that were relatively evenly matched for overall preference, but marmosets nevertheless very often show a mild preference. Consequently, we found that marmosets tended to be more willing to continue working for a juice that was different from the free juice if the response-contingent juice was their preferred juice. Therefore, for all manipulations where a marmoset showed a mild preference between juices, the preferred juice was always assigned to be the response-contingent juice in the non-degraded sessions, and the non-preferred juice was always the response-contingent juice in the degraded sessions (which is also the “free” juice). Because our measure of contingency degradation compares responding for a given juice in the degradation probe session (e.g. strawberry, in the presence of “free” juice) with its control session (e.g. strawberry, in the absence of “free juice”), all within a block, any slight juice preference will not influence any contingency degradation effect observed.

In addition to the four-day contingency degradation block, a four-day baseline block was also conducted, which consisted of four control sessions (Figure S1). Marmosets received intracranial infusions on the final two sessions of the block, in which in one session they respond to receive Juice 1, the response-contingent reward used in the degraded sessions of the contingency degradation task, and the other session to Juice 2, the response-contingent reward used in the non-degraded sessions of the contingency degradation task. These control sessions enabled determination of the manipulation’s effects on baseline responding for reward, separate from any effects on responding mediated by changes in response contingencies. Thus, no free rewards were given in baseline sessions. Behavior measure was calculated the same way as for the degradation sessions, without, of course, the need to account for free rewards (see below).

For the prefrontal and cingulate brain regions there were three manipulations (saline, inactivation via muscimol/baclofen, over-activation via DHK) and for the caudate nucleus, there were two manipulations (saline and inactivation via CNQX). Whenever possible infusions were counterbalanced. Where a brain region was reached by extending the injector, the region above was always infused first, i.e. area 24 was infused before area 14-25 and area 32 was infused before area 14. Otherwise, where animals had cannulae in more than one brain region, infusions in brain regions were counterbalanced accordingly, i.e. area 11 was infused before or after area 24 and area 32 was infused before or after the caudate. Counterbalancing was also implemented with respect to whether (i) degradation sessions occurred before or after baseline and (ii) saline occurred before or after the experimental manipulation. Since the contingency degradation and baseline blocks consisted of 4 sessions, each block took place between Monday to Friday. Depending upon performance in the first two sessions of the block, in some weeks the marmosets just received control blocks with no infusions to ensure their performance was stable between infusion blocks.

##### Behavioral measures

The main behavioral measure was the contingency degradation index (CDI). This was calculated in the degradation sessions as follows:

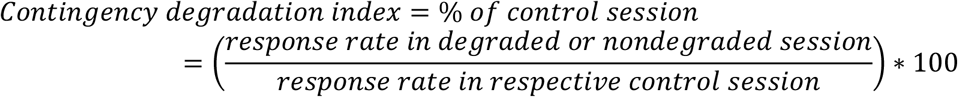

This approach, measuring response rate as a percentage of that of the same subject in a control session, accounts for animals’ individual variability in response rate.

In the baseline sessions, an equivalent CDI-like measure was also used:

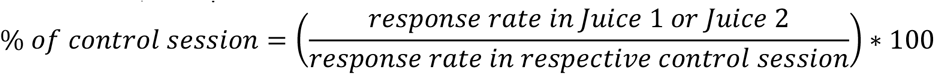

During reward collection periods, animals did not have access to the touchscreen stimulus for responding. To take into account the additional time animals spent drinking during degradation probe sessions with additional (free) reward, compared to control sessions, we calculated the index above to compare response rates during non-reward collection periods, rather than response numbers.

Response rate (responses per min.) in control sessions is calculated as follows:

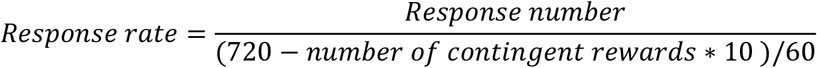

Where 720 is session length in seconds and 10 is reward duration in seconds. The response rate in degradation sessions:

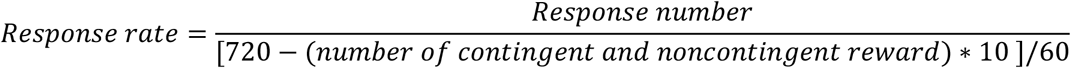

##### Cannulation procedure

Marmosets were premedicated with ketamine hydrochloride (Vetalar; 0.05 mL of a 100 mg/mL solution, i.m.; Amersham Biosciences and Upjohn, Crawley, UK) and then given a long-lasting nonsteroidal, anti-inflammatory analgesic (Carprieve; 0.03 mL of 50 mg/mL carprofen, s.c.; Pfizer, Kent, UK). They were intubated (using Intubeaze 20mg/ml lidocaine hydrochloride spray, Dechra Veterinary Products Ltd., Shropshire, UK), placed into a stereotaxic frame modified for the marmoset (David Kopf, Tujanga, CA) and maintained on 2.0–2.5% isoflurane in 0.3 L/min O_2_ throughout the surgery. Heart rate, O_2_ saturation, respiratory rate, and CO_2_ saturation were all monitored by pulse oximetry and capnography (Microcap Handheld Capnograph, Oridion Capnography Inc., MA, USA) while core body temperature was monitored rectally (TES-1319 K-type digital thermometer, TES Electrical Electronic Corp., Taipei, Taiwan). Cannulae (Plastics One) were lowered bilaterally into desired brain regions using the stereotaxic arm. The co-ordinates for each brain region are listed in Table S3, and brain implant locations for each animal in Table 1. Coordinates were adjusted in situ where necessary based on cortical depth within the prefrontal cortex at +17.5 anteroposterior (AP), −1.5 lateromedial (LM) as previously reported (Roberts et al., 2007); this adjustment varied between −0.5 and −1.0mm. An extra depth check was performed for area 11 at its target AP and LM coordinates to obtain the target depth from the cortex. Each animal received bilateral cannulae in two target regions, areas 24 and 11, and area 32 and caudate nucleus. Access to area 14-25 or area 14 was via extended injectors through cannulae (vertically placed) in area 24 or area 32, respectively. Postoperatively, monkeys received the analgesic meloxicam (0.1 mL of a 1.5 mg/mL oral suspension; Boehringer Ingelheim, Germany) for the next 3 days as well as at least a full 7 days of “weekend diet” and water *ad libitum* to ensure complete recovery before returning to testing. The implants were cleaned with 70% ethanol during every infusion and at least once every week (and caps and cannula dummies changed) to ensure the cannula site remained free from infection.

##### Intracerebral drug infusion

The infusions were conducted using aseptic procedures. The injectors were connected to 10 µL syringes (Hamilton), which were mounted on an infusion pump. The marmoset was held comfortably by a researcher, the dust caps and dummy cannulae were removed, the guide cannulae were cleaned with 70% ethanol wipes. The injectors were placed into the guide cannulae, extending 1.5mm below the cannulae for areas 32, 1.0mm for area 24, area 11 and the caudate nucleus, 3.5mm for area 14 and 4.5mm for area 14-25. Bilateral infusions were carried out; for more information on the drugs infused, please see Table S3. Injectors were left in place for one additional minute for drugs to diffuse. The injectors were then taken out, dummy and caps replaced on the guide cannulae, and the marmoset was returned to the home cage.

#### Post-mortem histological processing

##### Assessment of cannula placement

At the end of the experiment, all monkeys were sedated with ketamine hydrochloride (Pharmacia and Upjohn, 0.05 mL of a 100 mg/mL solution, i.m.) and humanely euthanized with Euthatal (1 mL of a 200 mg/mL solution, pentobarbital sodium; Merial Animal Health Ltd; i.v.) before being perfused transcardially with 400 mL of 0.1 M phosphate-buffered saline (PBS), followed by 400 mL of 4% paraformaldehyde fixative solution over approximately 15 minutes. The cannulae and dental cement were carefully removed. After the brain was removed, it was left in the 4% paraformaldehyde fixative solution overnight, before being transferred to 0.01M PBS-azide solution for at least 48 hours and then transferred to 30% sucrose solution for a further 48 hours for cryoprotection. Brains were sectioned on a freezing microtome (coronal sections; 40-60mm), mounted on slides and stained with cresyl violet. The sections were viewed under a Leitz DMRD microscope (Leica Microsystems, Wetzlar, Germany). The cannula locations for each animal were represented on schematized coronal sections of the marmoset brain (Figure 2). Before euthanasia, some animals underwent infusions of drugs for *c-fos* verification and one animal underwent an anatomical tract-tracing study.

##### Tract tracer infusion, immunohistochemistry protocol and image analysis

The left caudate nucleus of one monkey (not included in the behavioral study) was infused with the retrograde tracer cholera toxin B subunit (C9903, Sigma-Aldrich) via guide cannulae. The rate of infusion was 0.1 µL/min for 2 minutes, with 25 minutes of wait time for the drug to diffuse. The monkey was perfused after 10 days and the brain was processed and cut. Each section was 40 μm thick, and one in every five sections was taken for immunohistochemistry. On day 1, the brain sections were put into well plates to wash three times for 10 minutes each in 0.1M Tris-NaCl (pre-made the day before, pH adjusted to 7.4; Tris-base, T4661-100g, Sigma-Aldrich; NaCl – S7653-1Kg, Sigma-Aldrich). The washes occurred at room temperature and the wells were placed on a rocker. The 0.1M Tris-NaCl was changed between each wash in all situations. The sections were quenched to prevent endogenous peroxidase activity in 10% methanol and 10% H_2_O_2_ mixed solution for 5 minutes. The sections were then washed again three times for 10 minutes each in 0.1M Tris-NaCl. The sections were blocked in 0.1M Tris-NaCl with 0.2% Triton X-100 and 1% normal swine serum (S-4000, VectorLabs) for one hour at room temperature on a rocker. The sections were incubated overnight at room temperature, placed on a rocker, immersed in 0.1M Tris-NaCl with 0.2% Triton X-100, 1% normal swine serum and 1:2000 goat anti-choleragenoid primary antibody (703, Quadratech). On day 2, the brain sections were washed three times for 10 minutes each in 0.1M Tris-NaCl. They were then incubated for two hours at room temperature on a rocker, in 0.1M Tris-NaCl with 0.2% Triton X-100 and 1:200 biotinylated donkey anti-goat secondary antibody (bs-0294D-Biotin-BSS, Stratech). The brain sections were washed three times for 10 minutes each in 0.1M Tris-NaCl. They were incubated for 90 minutes at room temperature on a rocker with a ready-to-use avidin-biotin complex. The brain sections were washed three times for 10 minutes each in 0.1M Tris-NaCl. The sections were reacted with 3,3′-diaminobenzidine (DAB), using the ImmPactDAB horseradish peroxidase (HRP) Substrate Kit (SK-4100, Vector Labs). The reaction time inside DAB was determined empirically under the microscope. Once the desired staining was achieved, the section was immediately transferred to ice-cold 0.01M PBS to terminate the DAB reaction. The brain sections were mounted on gelatin-coated slides and dried overnight at room temperature. They were then dehydrated for 2 minutes each in solutions in the following order: 100% ddH_2_O, 25% ddH_2_O/75% ETOH, 100% ETOH, 50% ETOH/50% Xylene, 100% Xylene. The slides were coverslipped with DPX.

Images were acquired under bright field using a stereomicroscope (M205 FA; Leica, Wetzlar, Germany). Cell counting was conducted automatically using ilastik (version 1.3.3) (Berg et al., 2019) and FIJI (Schindelin et al., 2012).

### Quantification and Statistical Analysis

Data were analyzed using a mixed-model ANOVA using R version 3.5.1 (R Development Core Team, 2020). We used the lme4 package to conduct linear mixed-effects models with Type III analysis of variance with Satterthwaite’s method for degrees of freedom (Bates et al., 2015). Bartlett’s test was used to determine the homogeneity of variance. Each significant main effect (p<0.05) was further examined using pair-wise comparisons of least square means (lsmeans package in R) for specified factors in linear or mixed models. Fixed factors were the between-subject factor *infusion area* (region; area 11, area 24, area 14-25, area 32, area 14 and caudate nucleus) and the within-subject factors *treatment* (saline, mus/bac, DHK for PFC sub-regions; saline and CNQX for caudate nucleus) and *degradation* (degraded vs. non-degraded). *Subject* was a random factor. To account for individual variabilities in response rate, the dependent variable was the contingency degradation index. Data for areas 24, 11 and 14-25 on degradation sessions were square-root transformed to satisfy the assumptions of the analysis of variance but the data presented in graphs are not transformed for comparison purposes. Data from each individual brain region were analyzed separately. Data from drug manipulations on baseline sessions underwent the same analysis. We used the standard error of difference of the means (SED) as a more appropriate indication of the variance between means than the standard error of the mean (SEM), following ANOVA. The SED is calculated according to the equation given in Cochran and Cox (1957, p31).

Data from control and degradation sessions in the absence of manipulations (Figure 1D, E) were analyzed using within-subject repeated measures ANOVA in R (afex package; R Core Team, 2020). Factors for the response rate data (Figure 1D) include two within-group factors of *degradation* (degraded vs. non-degraded) and *free juice* (presence vs. absence). All graphs were first completed in GraphPad Prism version 8.3.0 for Windows (GraphPad Software, La Jolla, California, USA), then transferred to Adobe Illustrator CS6 (Adobe Inc., San Jose, California, USA) for aesthetics.

## Supplemental Figures

**Figure S1.**
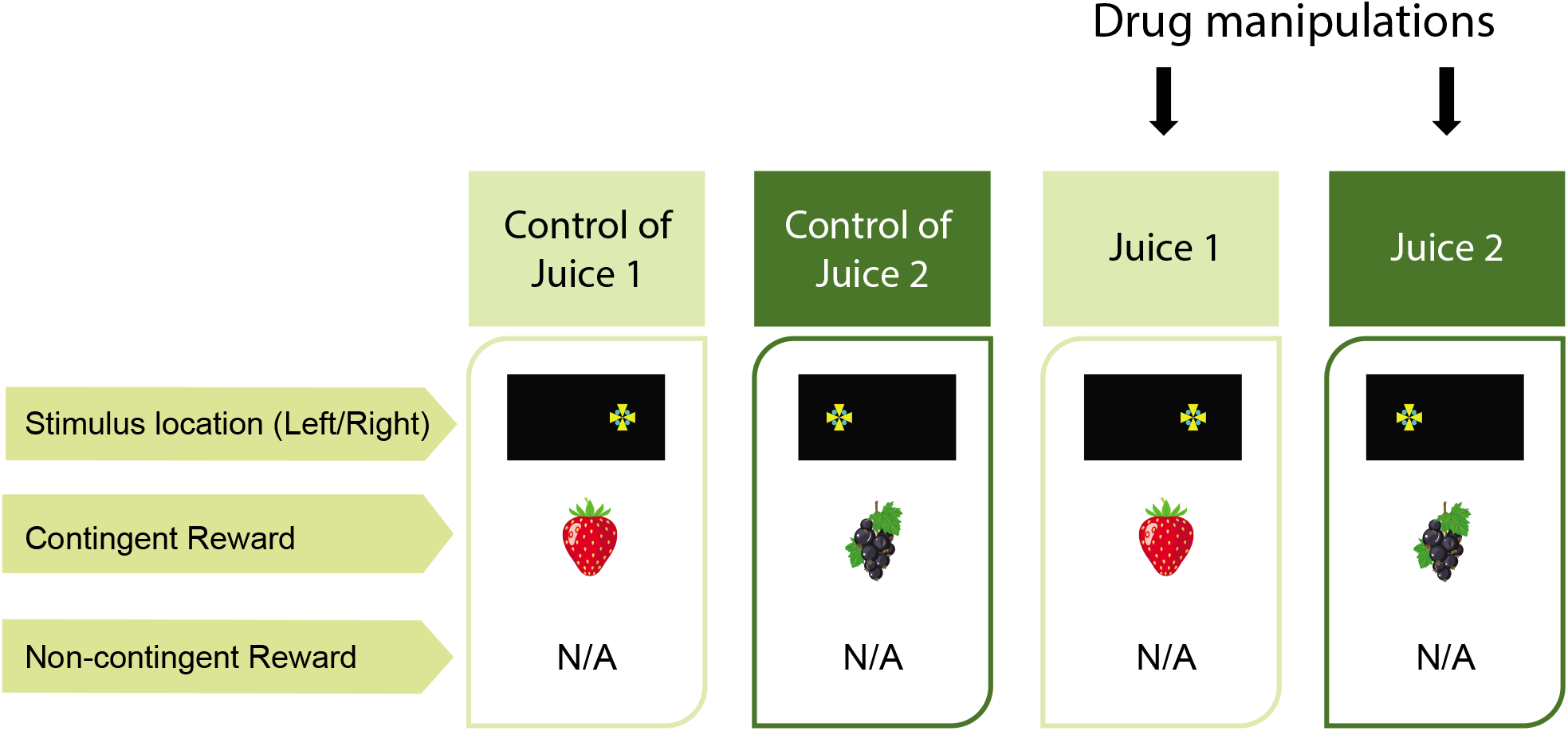
Task procedure for baseline sessions. Related to Figures 2, 5, S2 and Methods. Baseline sessions are four-day blocks. In the first two days, marmosets respond to two action-outcome (A-O) associations on separate days, in which one of the A-O associations is going to be degraded and the other to not be degraded in the degradation sessions. The last two days are the same as the first two days but with marmosets receiving drug manipulations prior to testing. No free, non-contingent reward was present in any conditions.

**Figure S2.**
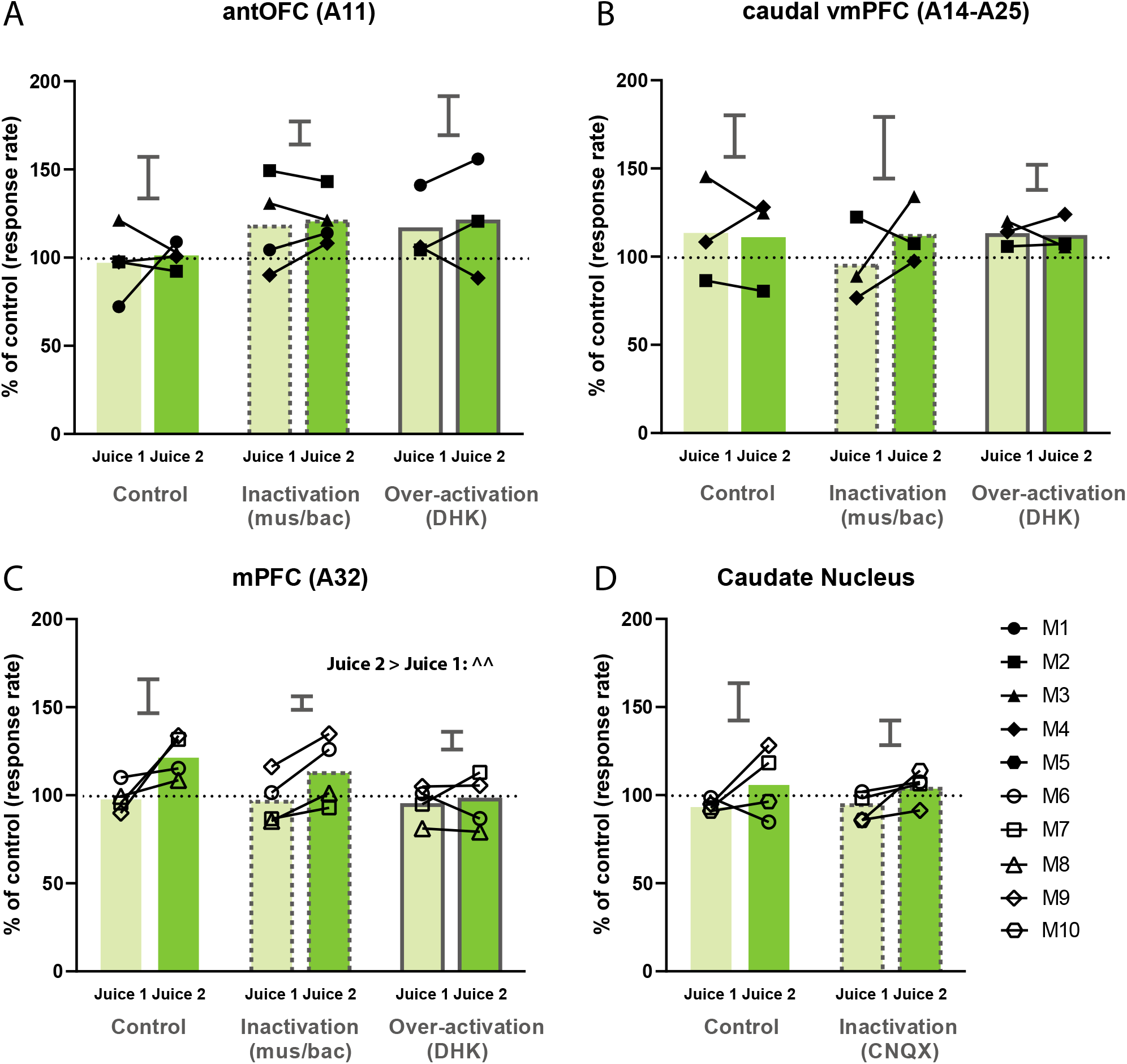
Effects of control, inactivation or over-activation of critical PFC and caudate nucleus regions in baseline sessions. Related to Figure 5. **(A, B)** Analysis of area 11 or area 14-25 baseline sessions revealed no main effects of juice conditions (area 11: F1, 13.069 = 0.209, p = 0.655; area 14-25: F1, 10 = 0.245, p = 0.632) or treatments (area 11: F2, 13,684 = 2.684, p = 0.104; area 14-25: F2, 10 = 0.324, p = 0.731). **(C)** For area 32, a main effect of juice conditions was observed (F1, 15 = 9.338, p = 0.00801), where marmosets significantly increased responding in Juice 2 when compared to Juice 1 across all drug manipulations (p = 0.008). **(D)** Analysis of caudate nucleus baseline sessions revealed no main effects of juice conditions (F1, 12 = 4.084, p = 0.0662) or treatments (F1, 12 = 0.0696, p = 0.796). Relevant graphs show 2 X SED for “Jucie 1 v. Juice 2” comparisons (area 11: n = 4; area 32: n = 4; area 14-25: 3). ^ indicates significant effect between juice conditions. ^^: p < 0.01

**Figure S3.**
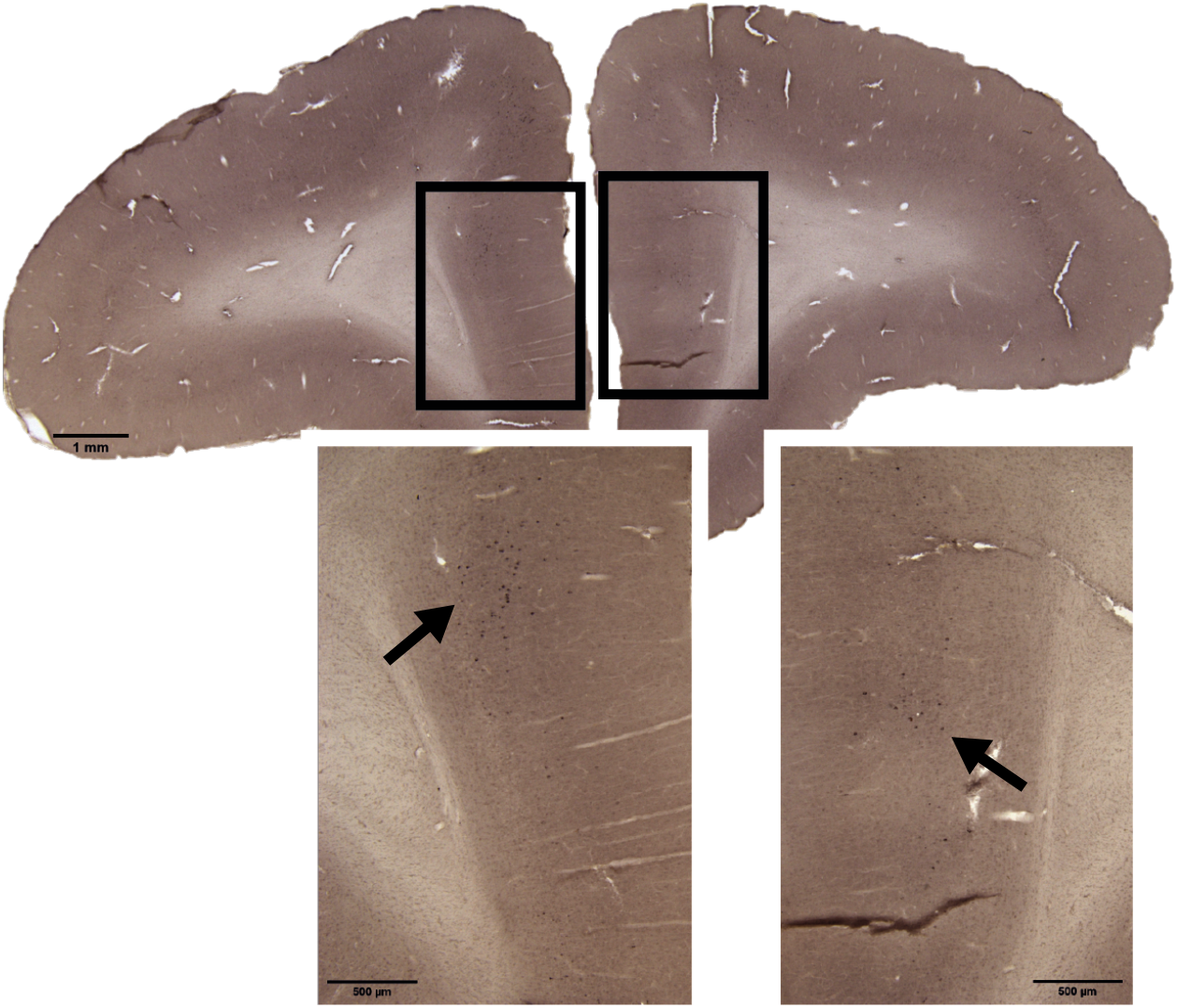
Area 32 targeted in this study also projects to the caudate nucleus region that receives area 24 projection. Related to Figure 4. The same retrograde tracer choleratoxin B subunit injection (left hemisphere) as Figure 4.

## Tables

**Table S1.** Touchscreen training schedule. Related to Figure 1 and Methods.

| Schedule | Training Phase | Stimulus | Juice | Reward Length | ITI |
| --- | --- | --- | --- | --- | --- |
| FT | 1 | green bar | Banana milkshake | 8 sec | 1 sec |
| FT | 2 | green square centre | Banana milkshake | 8 sec | 1 sec |
| FT | 3 | green square L/R side | Banana milkshake | 8 -> 5 sec | 1 -> 3sec |
| FR1 | 3 | green square L/R side | Banana milkshake | 5 sec | - |
| VR 3 | 3 | green square L/R side | Banana milkshake | 5 sec | - |
| VR 6 | 3 | green square L/R side | Banana milkshake | 7.5 sec | - |
| VR 10 | 3 | green square L/R side | Banana milkshake | 10 sec | - |
| VR 10 | 3 | green square L/R side | Juice | 10 sec | - |
| Contingency | 3 | green square L/R side | Juice | 10 sec | - |
| Contingency | 3 | Maltese cross L/R side | Juice | 10 sec | - |

**Table S2.** Cannulation co-ordinates. Related to Table 1. AP: anteroposterior; LM: lateromedial; *Area 14-25 and area 14 were reached by extending the injectors via the area 24 and area 32 guide cannulae, respectively. ^the caudate nucleus guide cannula was at 10 degrees angle away from the inter-aural line. Therefore, the LM of the guide entering the brain surface is +/-3.2mm, whereas the actual targeted location inside the caudate nucleus is +/-2.2mm and 5.0mm vertically from the brain surface.

| Area | AP co-ordinate (mm) | LM co-ordinate (mm) | Depth (mm) |
| --- | --- | --- | --- |
| Area 11 | +17.0 | +/- 3.0 | 1.7<br>(from base) |
| Area 24* | +15.4 | +/- 1.0 | 2.5<br>(from surface) |
| Area 32* | +16.8 | +/- 1.0 | 1.5<br>(from surface) |
| Caudate | +11.0 | +/- 2.2^ | 5.0^<br>(from surface) |

**Table S3.** Drugs used in the study. Related to Figures 3, 4, 5.

| Drug(s) | Mechanism | Concentration | Infusion Rate | Pre-Treatment Time | Source |
| --- | --- | --- | --- | --- | --- |
| Muscimol-Baclofen (Mus-Bac) | Muscimol: GABA<sub>A</sub> receptor antagonist<br>Baclofen: GABA<sub>B</sub> receptor antagonist | Muscimol: 0.1mM<br>Baclofen: 1.0mM | 0.25µl/min for 2 mins | 25 minutes | Sigma-Aldrich, St Louis, USA |
| Dihydrokainic acid (DHK) | Excitatory amino acid transporter-2 (EAAT2/GLT-1) inhibitor | 6.25 nmol/µL | 0.50µl/min for 2 mins | 8-15 minutes | Tocris, Bristol, UK |
| CNQX | selective AMPA/Kainate receptor antagonist | 1.0mM | 0.3 µL/min for 1 min | 8 minutes | Tocris, Bristol, UK |

## References

Alexander, L., Gaskin, P.L.R., Sawiak, S.J., Fryer, T.D., Hong, Y.T., Cockcroft, G.J., Clarke, H.F., and Roberts, A.C. (2019). Fractionating Blunted Reward Processing Characteristic of Anhedonia by Over-Activating Primate Subgenual Anterior Cingulate Cortex. Neuron 101, 307–320 e306.

Amiez, C., Joseph, J.P., and Procyk, E. (2006). Reward encoding in the monkey anterior cingulate cortex. Cereb Cortex 16, 1040–1055.

Anderson, C.M., and Swanson, R.A. (2000). Astrocyte glutamate transport: review of properties, regulation, and physiological functions. Glia 32, 1–14.

Arriza, J.L., Fairman, W.A., Wadiche, J.I., Murdoch, G.H., Kavanaugh, M.P., and Amara, S.G. (1994). Functional comparisons of three glutamate transporter subtypes cloned from human motor cortex. J Neurosci 14, 5559–5569.

Averbeck, B.B., Lehman, J., Jacobson, M., and Haber, S.N. (2014). Estimates of projection overlap and zones of convergence within frontal-striatal circuits. J Neurosci 34, 9497–9505.

Balleine, B.W. (2019). The Meaning of Behavior: Discriminating Reflex and Volition in the Brain. Neuron 104, 47–62.

Balleine, B.W., and Dickinson, A. (1998). Goal-directed instrumental action: contingency and incentive learning and their cortical substrates. Neuropharmacology 37, 407–419.

Balleine, B.W., Leung, B.K., and Ostlund, S.B. (2011). The orbitofrontal cortex, predicted value, and choice. Ann N Y Acad Sci 1239, 43–50.

Balleine, B.W., and O’Doherty, J.P. (2010). Human and rodent homologies in action control: corticostriatal determinants of goal-directed and habitual action. Neuropsychopharmacology 35, 48–69.

Barch, D.M., and Dowd, E.C. (2010). Goal representations and motivational drive in schizophrenia: the role of prefrontal-striatal interactions. Schizophrenia bulletin 36, 919–934.

Bates, D., Mächler, M., Bolker, B., and Walker, S. (2015). Fitting Linear Mixed-Effects Models Using lme4. Journal of Statistical Software; Vol 1, Issue 1 (2015).

Baxter, L.R., Jr., Schwartz, J.M., Mazziotta, J.C., Phelps, M.E., Pahl, J.J., Guze, B.H., and Fairbanks, L. (1988). Cerebral glucose metabolic rates in nondepressed patients with obsessive-compulsive disorder. Am J Psychiatry 145, 1560–1563.

Bechtholt-Gompf, A.J., Walther, H.V., Adams, M.A., Carlezon, W.A., Jr., Ongur, D., and Cohen, B.M. (2010). Blockade of astrocytic glutamate uptake in rats induces signs of anhedonia and impaired spatial memory. Neuropsychopharmacology 35, 2049–2059.

Berg, S., Kutra, D., Kroeger, T., Straehle, C.N., Kausler, B.X., Haubold, C., Schiegg, M., Ales, J., Beier, T., Rudy, M., et al. (2019). ilastik: interactive machine learning for (bio)image analysis. Nat Methods 16, 1226–1232.

Bradfield, L., Morris, R., and Balleine, B. (2017). Obsessive-compulsive disorder as a failure to integrate goal-directed and habitual action control. p. 1872.

Bradfield, L.A., Dezfouli, A., van Holstein, M., Chieng, B., and Balleine, B.W. (2015). Medial Orbitofrontal Cortex Mediates Outcome Retrieval in Partially Observable Task Situations. Neuron 88, 1268–1280.

Bradfield, L.A., Hart, G., and Balleine, B.W. (2018). Inferring action-dependent outcome representations depends on anterior but not posterior medial orbitofrontal cortex. Neurobiology of learning and memory 155, 463–473.

Burman, K.J., and Rosa, M.G. (2009). Architectural subdivisions of medial and orbital frontal cortices in the marmoset monkey (Callithrix jacchus). J Comp Neurol 514, 11–29.

Cardinal, R.N., and Aitken, M.R.F. (2010). Whisker: A client—server high-performance multimedia research control system. Behavior Research Methods 42, 1059–1071.

Cartoni, E., Balleine, B., and Baldassarre, G. (2016). Appetitive Pavlovian-instrumental Transfer: A review. Neuroscience & Biobehavioral Reviews 71, 829–848.

Choi, E.Y., Tanimura, Y., Vage, P.R., Yates, E.H., and Haber, S.N. (2017). Convergence of prefrontal and parietal anatomical projections in a connectional hub in the striatum. Neuroimage 146, 821–832.

Chudasama, Y., Daniels, T.E., Gorrin, D.P., Rhodes, S.E., Rudebeck, P.H., and Murray, E.A. (2013). The role of the anterior cingulate cortex in choices based on reward value and reward contingency. Cereb Cortex 23, 2884–2898.

Cochran, W.G., and Cox, G.M. (1957). Experimental Designs (2nd edition, New York: Wiley), pp. 31.

Corbit, L.H., and Balleine, B.W. (2003). The role of prelimbic cortex in instrumental conditioning. Behav Brain Res 146, 145–157.

Corbit, L.H., and Janak, P.H. (2010). Posterior dorsomedial striatum is critical for both selective instrumental and Pavlovian reward learning. The European journal of neuroscience 31, 1312–1321.

de Jonge, J.C., Vinkers, C.H., Hulshoff Pol, H.E., and Marsman, A. (2017). GABAergic Mechanisms in Schizophrenia: Linking Postmortem and In Vivo Studies. Frontiers in Psychiatry 8.

de Wit, S., and Dickinson, A. (2009). Associative theories of goal-directed behaviour: a case for animal-human translational models. Psychol Res 73, 463–476.

de Wit, S., Kindt, M., Knot, S.L., Verhoeven, A.A.C., Robbins, T.W., Gasull-Camos, J., Evans, M., Mirza, H., and Gillan, C.M. (2018). Shifting the balance between goals and habits: Five failures in experimental habit induction. J Exp Psychol Gen 147, 1043–1065.

Dickinson, A., and Weiskrantz, L. (1985). Actions and habits: the development of behavioural autonomy. Philosophical Transactions of the Royal Society of London B, Biological Sciences 308, 67–78.

Dolan, R.J., and Dayan, P. (2013). Goals and habits in the brain. Neuron 80, 312–325.

Draganski, B., Kherif, F., Kloppel, S., Cook, P.A., Alexander, D.C., Parker, G.J., Deichmann, R., Ashburner, J., and Frackowiak, R.S. (2008). Evidence for segregated and integrative connectivity patterns in the human Basal Ganglia. J Neurosci 28, 7143–7152.

Ersche, K.D., Gillan, C.M., Jones, P.S., Williams, G.B., Ward, L.H., Luijten, M., de Wit, S., Sahakian, B.J., Bullmore, E.T., and Robbins, T.W. (2016). Carrots and sticks fail to change behavior in cocaine addiction. Science 352, 1468–1471.

Everitt, B.J., and Robbins, T.W. (2005). Neural systems of reinforcement for drug addiction: from actions to habits to compulsion. Nat Neurosci 8, 1481–1489.

Fallgren, A.B., and Paulsen, R.E. (1996). A microdialysis study in rat brain of dihydrokainate, a glutamate uptake inhibitor. Neurochem Res 21, 19–25.

Fitzgerald, K.D., Welsh, R.C., Stern, E.R., Angstadt, M., Hanna, G.L., Abelson, J.L., and Taylor, S.F. (2011). Developmental alterations of frontal-striatal-thalamic connectivity in obsessive-compulsive disorder. J Am Acad Child Adolesc Psychiatry 50, 938–948 e933.

Gillan, C.M., Morein-Zamir, S., Kaser, M., Fineberg, N.A., Sule, A., Sahakian, B.J., Cardinal, R.N., and Robbins, T.W. (2014). Counterfactual processing of economic action-outcome alternatives in obsessive-compulsive disorder: further evidence of impaired goal-directed behavior. Biol Psychiatry 75, 639–646.

Gillan, C.M., and Robbins, T.W. (2014). Goal-directed learning and obsessive-compulsive disorder. Philos Trans R Soc Lond B Biol Sci 369.

Gremel, C.M., and Costa, R.M. (2013). Orbitofrontal and striatal circuits dynamically encode the shift between goal-directed and habitual actions. Nature Communications 4, 2264.

Haber, S.N. (2003). The primate basal ganglia: parallel and integrative networks. Journal of Chemical Neuroanatomy 26, 317–330.

Hammond, L.J. (1980). The effect of contingency upon the appetitive conditioning of free-operant behavior. J Exp Anal Behav 34, 297–304.

Hart, G., Bradfield, L.A., and Balleine, B.W. (2018a). Prefrontal Corticostriatal Disconnection Blocks the Acquisition of Goal-Directed Action. J Neurosci 38, 1311–1322.

Hart, G., Bradfield, L.A., Fok, S.Y., Chieng, B., and Balleine, B.W. (2018b). The Bilateral Prefronto-striatal Pathway Is Necessary for Learning New Goal-Directed Actions. Current biology : CB 28, 2218-2229.e2217.

Hayden, B.Y., Heilbronner, S.R., Pearson, J.M., and Platt, M.L. (2011). Surprise signals in anterior cingulate cortex: neuronal encoding of unsigned reward prediction errors driving adjustment in behavior. J Neurosci 31, 4178–4187.

Hayden, B.Y., Pearson, J.M., and Platt, M.L. (2009). Fictive reward signals in the anterior cingulate cortex. Science 324, 948–950.

Hayden, B.Y., and Platt, M.L. (2010). Neurons in anterior cingulate cortex multiplex information about reward and action. The Journal of neuroscience : the official journal of the Society for Neuroscience 30, 3339–3346.

Heilbronner, S.R., Rodriguez-Romaguera, J., Quirk, G.J., Groenewegen, H.J., and Haber, S.N. (2016). Circuit-Based Corticostriatal Homologies Between Rat and Primate. Biol Psychiatry 80, 509–521.

Heyes, C., and Dickinson, A. (1990). The Intentionality of Animal Action. Mind & Language 5, 87–103.

Hiser, J., and Koenigs, M. (2018). The Multifaceted Role of the Ventromedial Prefrontal Cortex in Emotion, Decision Making, Social Cognition, and Psychopathology. Biol Psychiatry 83, 638–647.

Holmes, N.M., Marchand, A.R., and Coutureau, E. (2010). Pavlovian to instrumental transfer: A neurobehavioural perspective. Neuroscience & Biobehavioral Reviews 34, 1277–1295.

Holroyd, C.B., and Coles, M.G.H. (2002). The neural basis of human error processing: reinforcement learning, dopamine, and the error-related negativity. Psychol Rev 109, 679–709.

Holroyd, C.B., and Yeung, N. (2012). Motivation of extended behaviors by anterior cingulate cortex. Trends Cogn Sci 16, 122–128.

Jackson, S.A., Horst, N.K., Pears, A., Robbins, T.W., and Roberts, A.C. (2016). Role of the Perigenual Anterior Cingulate and Orbitofrontal Cortex in Contingency Learning in the Marmoset. Cereb Cortex 26, 3273–3284.

Kennerley, S.W., Walton, M.E., Behrens, T.E., Buckley, M.J., and Rushworth, M.F. (2006). Optimal decision making and the anterior cingulate cortex. Nat Neurosci 9, 940–947.

Laubach, M., Amarante, L.M., Swanson, K., and White, S.R. (2018). What, If Anything, Is Rodent Prefrontal Cortex? eneuro 5, ENEURO.0315-0318.2018.

Liljeholm, M., Tricomi, E., O’Doherty, J.P., and Balleine, B.W. (2011). Neural correlates of instrumental contingency learning: differential effects of action-reward conjunction and disjunction. J Neurosci 31, 2474–2480.

Mackey, S., and Petrides, M. (2010). Quantitative demonstration of comparable architectonic areas within the ventromedial and lateral orbital frontal cortex in the human and the macaque monkey brains. Eur J Neurosci 32, 1940–1950.

Maia, T.V., Cooney, R.E., and Peterson, B.S. (2008). The neural bases of obsessive-compulsive disorder in children and adults. Dev Psychopathol 20, 1251–1283.

Mailly, P., Aliane, V., Groenewegen, H.J., Haber, S.N., and Deniau, J.-M. (2013). The Rat Prefrontostriatal System Analyzed in 3D: Evidence for Multiple Interacting Functional Units. The Journal of Neuroscience 33, 5718–5727.

Majka, P., Bai, S., Bakola, S., Bednarek, S., Chan, J.M., Jermakow, N., Passarelli, L., Reser, D.H., Theodoni, P., Worthy, K.H., et al. (2020). Open access resource for cellular-resolution analyses of corticocortical connectivity in the marmoset monkey. Nat Commun 11, 1133.

Matsumoto, M., Matsumoto, K., Abe, H., and Tanaka, K. (2007). Medial prefrontal cell activity signaling prediction errors of action values. Nature Neuroscience 10, 647–656.

Menzies, L., Chamberlain, S.R., Laird, A.R., Thelen, S.M., Sahakian, B.J., and Bullmore, E.T. (2008). Integrating evidence from neuroimaging and neuropsychological studies of obsessive-compulsive disorder: the orbitofronto-striatal model revisited. Neurosci Biobehav Rev 32, 525–549.

Milad, M., Quirk, G., Roger, P., Orr, S., Fischl, B., and Rauch, S. (2007). A Role for the Human Dorsal Anterior Cingulate Cortex in Fear Expression. Biological psychiatry 62, 1191–1194.

Morris, R.W., Cyrzon, C., Green, M.J., Le Pelley, M.E., and Balleine, B.W. (2018). Impairments in action–outcome learning in schizophrenia. Translational Psychiatry 8, 54.

Morris, R.W., Quail, S., Griffiths, K.R., Green, M.J., and Balleine, B.W. (2015). Corticostriatal Control of Goal-Directed Action Is Impaired in Schizophrenia. Biological Psychiatry 77, 187–195.

Munoz, M.D., Herreras, O., Herranz, A.S., Solis, J.M., Martin del Rio, R., and Lerma, J. (1987). Effects of dihydrokainic acid on extracellular amino acids and neuronal excitability in the in vivo rat hippocampus. Neuropharmacology 26, 1–8.

Murray, E.A., O’Doherty, J.P., and Schoenbaum, G. (2007). What We Know and Do Not Know about the Functions of the Orbitofrontal Cortex after 20 Years of Cross-Species Studies. The Journal of Neuroscience 27, 8166–8169.

Nakao, T., Nakagawa, A., Yoshiura, T., Nakatani, E., Nabeyama, M., Yoshizato, C., Kudoh, A., Tada, K., Yoshioka, K., Kawamoto, M., et al. (2005). Brain activation of patients with obsessive-compulsive disorder during neuropsychological and symptom provocation tasks before and after symptom improvement: a functional magnetic resonance imaging study. Biol Psychiatry 57, 901–910.

Noonan, M.P., Walton, M.E., Behrens, T.E., Sallet, J., Buckley, M.J., and Rushworth, M.F. (2010). Separate value comparison and learning mechanisms in macaque medial and lateral orbitofrontal cortex. Proc Natl Acad Sci U S A 107, 20547–20552.

O’Callaghan, C., Vaghi, M.M., Brummerloh, B., Cardinal, R.N., and Robbins, T.W. (2019). Impaired awareness of action-outcome contingency and causality during healthy ageing and following ventromedial prefrontal cortex lesions. Neuropsychologia 128, 282–289.

O’Doherty, J.P., Cockburn, J., and Pauli, W.M. (2017). Learning, Reward, and Decision Making. Annual Review of Psychology 68, 73–100.

Ostlund, S.B., and Balleine, B.W. (2005). Lesions of medial prefrontal cortex disrupt the acquisition but not the expression of goal-directed learning. J Neurosci 25, 7763–7770.

Ostlund, S.B., and Balleine, B.W. (2007). Orbitofrontal cortex mediates outcome encoding in Pavlovian but not instrumental conditioning. J Neurosci 27, 4819–4825.

Padoa-Schioppa, C., and Assad, J.A. (2006). Neurons in the orbitofrontal cortex encode economic value. Nature 441, 223–226.

Panayi, M.C., and Killcross, S. (2018). Functional heterogeneity within the rodent lateral orbitofrontal cortex dissociates outcome devaluation and reversal learning deficits. eLife 7, e37357.

Parkes, S.L., Ravassard, P.M., Cerpa, J.-C., Wolff, M., Ferreira, G., and Coutureau, E. (2017). Insular and Ventrolateral Orbitofrontal Cortices Differentially Contribute to Goal-Directed Behavior in Rodents. Cerebral Cortex 28, 2313–2325.

Pauls, D.L., Abramovitch, A., Rauch, S.L., and Geller, D.A. (2014). Obsessive-compulsive disorder: an integrative genetic and neurobiological perspective. Nat Rev Neurosci 15, 410–424.

Paxinos, G., Watson, C., Petrides, M., Rosa, M., and Tokuno, H. (2012). The Marmoset Brain in Stereotaxic Coordinates (London: Elsevier Academic Press).

Price, J.L. (2007). Definition of the orbital cortex in relation to specific connections with limbic and visceral structures and other cortical regions. Ann N Y Acad Sci 1121, 54–71.

R Development Core Team (2020). R: A language and environment for statistical computing. (R Foundation for Statistical Computing).

Rauch, S.L., Jenike, M.A., Alpert, N.M., Bafter, L., Breiter, H.C.R., Savage, C.R., and Fischman, A.J. (1994). Regional Cerebral Blood Flow Measured During Symptom Provocation in Obsessive-Compulsive Disorder Using Oxygen 15— Labeled Carbon Dioxide and Positron Emission Tomography. Archives of General Psychiatry 51, 62–70.

Reber, J., Feinstein, J.S., O’Doherty, J.P., Liljeholm, M., Adolphs, R., and Tranel, D. (2017). Selective impairment of goal-directed decision-making following lesions to the human ventromedial prefrontal cortex. Brain 140, 1743–1756.

Rescorla, R.A. (1966). Predictability and number of pairings in Pavlovian fear conditioning. Psychonomic Science 4, 383–384.

Robbins, T.W., and Costa, R.M. (2017). Habits. Current biology : CB 27, R1200–r1206.

Robbins, T.W., Vaghi, M.M., and Banca, P. (2019). Obsessive-Compulsive Disorder: Puzzles and Prospects. Neuron 102, 27–47.

Roberts, A.C. (2020). Prefrontal Regulation of Threat-Elicited Behaviors: A Pathway to Translation. Annual Review of Psychology 71, 357–387.

Roberts, A.C., and Clarke, H.F. (2019). Why we need nonhuman primates to study the role of ventromedial prefrontal cortex in the regulation of threat- and reward-elicited responses. Proceedings of the National Academy of Sciences 116, 26297–26304.

Roberts, A.C., Tomic, D.L., Parkinson, C.H., Roeling, T.A., Cutter, D.J., Robbins, T.W., and Everitt, B.J. (2007). Forebrain connectivity of the prefrontal cortex in the marmoset monkey (Callithrix jacchus): an anterograde and retrograde tract-tracing study. J Comp Neurol 502, 86–112.

Rolls, E.T. (2004). The functions of the orbitofrontal cortex. Brain Cogn 55, 11–29.

Rudebeck, P.H., Behrens, T.E., Kennerley, S.W., Baxter, M.G., Buckley, M.J., Walton, M.E., and Rushworth, M.F. (2008). Frontal cortex subregions play distinct roles in choices between actions and stimuli. J Neurosci 28, 13775–13785.

Rudebeck, P.H., and Murray, E.A. (2011). Dissociable effects of subtotal lesions within the macaque orbital prefrontal cortex on reward-guided behavior. J Neurosci 31, 10569–10578.

Rushworth, M.F., and Behrens, T.E. (2008). Choice, uncertainty and value in prefrontal and cingulate cortex. Nat Neurosci 11, 389–397.

Rushworth, M.F., Behrens, T.E., Rudebeck, P.H., and Walton, M.E. (2007). Contrasting roles for cingulate and orbitofrontal cortex in decisions and social behaviour. Trends Cogn Sci 11, 168–176.

Rushworth, M.F., Noonan, M.P., Boorman, E.D., Walton, M.E., and Behrens, T.E. (2011). Frontal cortex and reward-guided learning and decision-making. Neuron 70, 1054–1069.

Rushworth, M.F., Walton, M.E., Kennerley, S.W., and Bannerman, D.M. (2004). Action sets and decisions in the medial frontal cortex. Trends Cogn Sci 8, 410–417.

Russchen, F.T., Bakst, I., Amaral, D.G., and Price, J.L. (1985). The amygdalostriatal projections in the monkey. An anterograde tracing study. Brain Research 329, 241–257.

Schindelin, J., Arganda-Carreras, I., Frise, E., Kaynig, V., Longair, M., Pietzsch, T., Preibisch, S., Rueden, C., Saalfeld, S., Schmid, B., et al. (2012). Fiji: an open-source platform for biological-image analysis. Nat Methods 9, 676–682.

Schneider, B., and Koenigs, M. (2017). Human lesion studies of ventromedial prefrontal cortex. Neuropsychologia 107.

Seo, H., and Lee, D. (2007). Temporal filtering of reward signals in the dorsal anterior cingulate cortex during a mixed-strategy game. J Neurosci 27, 8366–8377.

Shipman, M.L., Johnson, G.C., Bouton, M.E., and Green, J.T. (2019). Chemogenetic Silencing of Prelimbic Cortex to Anterior Dorsomedial Striatum Projection Attenuates Operant Responding. eneuro 6, ENEURO.0125-0119.2019.

Stalnaker, T.A., Cooch, N.K., and Schoenbaum, G. (2015). What the orbitofrontal cortex does not do. Nat Neurosci 18, 620–627.

Stawicka, Z.M., Massoudi, R., Horst, N.K., Koda, K., Gaskin, P.L.R., Alexander, L., Santangelo, A.M., McIver, L., Cockcroft, G.J., Wood, C.M., and Roberts, A.C. (2020). Ventromedial prefrontal area 14 provides opposing regulation of threat and reward-elicited responses in the common marmoset. Proceedings of the National Academy of Sciences 117, 25116–25127.

Tanaka, S.C., Balleine, B.W., and O’Doherty, J.P. (2008). Calculating consequences: brain systems that encode the causal effects of actions. J Neurosci 28, 6750–6755.

Tang, W., Jbabdi, S., Zhu, Z., Cottaar, M., Grisot, G., Lehman, J.F., Yendiki, A., and Haber, S.N. (2019). A connectional hub in the rostral anterior cingulate cortex links areas of emotion and cognitive control. Elife 8.

Tran-Tu-Yen, D.A., Marchand, A.R., Pape, J.R., Di Scala, G., and Coutureau, E. (2009). Transient role of the rat prelimbic cortex in goal-directed behaviour. Eur J Neurosci 30, 464–471.

Tricomi, E.M., Delgado, M.R., and Fiez, J.A. (2004). Modulation of caudate activity by action contingency. Neuron 41, 281–292.

Vaghi, M.M., Cardinal, R.N., Apergis-Schoute, A.M., Fineberg, N.A., Sule, A., and Robbins, T.W. (2019). Action-Outcome Knowledge Dissociates From Behavior in Obsessive-Compulsive Disorder Following Contingency Degradation. Biol Psychiatry Cogn Neurosci Neuroimaging 4, 200–209.

Valentin, V.V., Dickinson, A., and O’Doherty, J.P. (2007). Determining the neural substrates of goal-directed learning in the human brain. J Neurosci 27, 4019–4026.

Verhoeven, A., and de Wit, S. (2018). The Role of Habits in Maladaptive Behaviour and Therapeutic Interventions: Theory, Mechanisms, Change, and Contexts. pp. 285–303.

Vogt, B.A., Hof, P.R., Zilles, K., Vogt, L.J., Herold, C., and Palomero-Gallagher, N. (2013). Cingulate area 32 homologies in mouse, rat, macaque and human: cytoarchitecture and receptor architecture. J Comp Neurol 521, 4189–4204.

Wallis, C.U., Cardinal, R.N., Alexander, L., Roberts, A.C., and Clarke, H.F. (2017). Opposing roles of primate areas 25 and 32 and their putative rodent homologs in the regulation of negative emotion. Proc Natl Acad Sci U S A 114, E4075–E4084.

Wallis, J.D., and Kennerley, S.W. (2011). Contrasting reward signals in the orbitofrontal cortex and anterior cingulate cortex. Ann N Y Acad Sci 1239, 33–42.

Walton, M.E., Behrens, T.E., Buckley, M.J., Rudebeck, P.H., and Rushworth, M.F. (2010). Separable learning systems in the macaque brain and the role of orbitofrontal cortex in contingent learning. Neuron 65, 927–939.

Whiteside, S.P., Port, J.D., and Abramowitz, J.S. (2004). A meta-analysis of functional neuroimaging in obsessive-compulsive disorder. Psychiatry Res 132, 69–79.

Yin, H.H., Ostlund, S.B., Knowlton, B.J., and Balleine, B.W. (2005). The role of the dorsomedial striatum in instrumental conditioning. Eur J Neurosci 22, 513–523.

Zimmermann, K.S., Yamin, J.A., Rainnie, D.G., Ressler, K.J., and Gourley, S.L. (2017). Connections of the Mouse Orbitofrontal Cortex and Regulation of Goal-Directed Action Selection by Brain-Derived Neurotrophic Factor. Biol Psychiatry 81, 366–377.

